# Benchmarking Docking Protocols for GPCR Allosteric Modulators

**DOI:** 10.64898/2026.08.12.744492

**Authors:** Tyler D. Thompson, Yinglong Miao

## Abstract

G protein-coupled receptor (GPCR) allosteric modulators (AMs) offer significant therapeutic advantages over orthosteric drugs, yet structure-based virtual screening lacks validated protocols accounting for the conformational complexity of GPCR allosteric sites. We benchmark docking protocols using PDB experimental structures and structural ensembles derived from Gaussian accelerated Molecular Dynamics (GaMD) simulations across four Class A GPCRs (including the muscarinic M2 and M4 receptors, the β2-adrenergic receptor, and the C-C chemokine receptor type 2) with four programs (Glide HTVS, AutoDock Vina, DOCK3.8, and Boltz-2) against experimentally validated modulator libraries and property-matched decoys. GaMD ensemble docking improved early AM enrichment across all four targets under at least one program. Glide ensemble docking was the only protocol to consistently improve early AM recovery across all four targets, ranking known actives almost exclusively within the top 0.5% of compounds at CCR2 and improving M2R active recovery nearly 9-fold relative to the PDB structure. GaMD free-energy landscape topology governed ensemble re-ranking strategy selection: population-skewed landscapes favored top binding energy ranking (*BE_min_*) while flat, multi-populated landscapes favored average binding energy ranking (*BE_avg_*), and at targets with dominant low-energy states, a single GaMD cluster matched or exceeded full ensemble or PDB performance. Taking the union of top percentile hits identified by both ensemble re-ranking methods, *BE_min_* / *BE_avg_*, maximizes chemical diversity at the earliest percentiles. Program-specific scaffold recovery biases further motivated a consensus *BE_min_* / *BE_avg_* approach to maximize hit diversity. The Boltz-2 deep-learning program showed minimal sensitivity to GaMD templates and underperformed conventional docking, suggesting its affinity predictions complement rather than replace physics- and empirical-based docking approaches for GPCR AM screening.

## INTRODUCTION

G protein-coupled receptors (GPCRs), the largest family of transmembrane proteins (>800 members) in the human genome, are implicated in a diverse range of therapeutic indications, including neurological, pulmonary, endocrine, and cardiac disorders^1,2^. Approximately 36% of all approved therapeutics target these receptors^2^. The vast majority of these drugs (∼92%) bind to the endogenous ligand-binding site (“orthosteric”), inducing conformational changes in the GPCR complex that directly affect downstream signaling^3^. However, because this pocket is highly conserved across GPCR subtypes, orthosteric ligands often lack receptor-subtype selectivity, limiting the potential for pharmacological interventions and contributing to deleterious off-target effects.^4,5^ Allosteric modulators (AMs) circumvent this limitation by binding to sites on the GPCR that are topologically distinct and often less conserved.^1,6–8^ These compounds can remotely shift the receptor’s conformational equilibrium toward either active or inactive states. Beyond their selectivity advantages, AMs exhibit remarkable functional diversity: positive AMs (PAMs) enhance orthosteric ligand affinity and/or efficacy, while negative AMs (NAMs) diminish it, with some AMs additionally exhibiting intrinsic efficacy or selective transducer bias. Despite their therapeutic promise, only ∼1% of GPCR-targeting drugs are FDA-approved AMs,^3^ largely due to the absence of robust structure-based drug design (SBDD) approaches that adequately account for the dynamic, conformation-dependent mechanisms underlying AM action at the GPCR.^9^

Traditional SBDD leverages static, 3D structural representations of biomolecular complexes derived from X-ray crystallography or cryo-EM, ideally in complex with known binders.^10^ For GPCR modulator discovery, however, this approach reaches a structural bottleneck. The AlloSteric Database (ASD) 2023 reports over 40,000 allosteric modulators across 200 unique GPCRs, yet only ∼100 structures in the GPCRdb contain a bound allosteric ligand, severely limiting the availability of structurally characterized allosteric sites.^2,11,12^ In the absence of experimental structures, deep learning (DL) co-folding methods (e.g., AlphaFold3, Boltz-2, Chai-1) have demonstrated great promise in predicting high-confidence protein-ligand complexes.^13–15^ However, their predictive power for GPCR allosteric binding remains poorly characterized. Even with a suitable starting structure, allosteric binding involves conformational fluctuations in pocket side-chains and loop regions unaccounted for in any single rigid structure, limiting downstream VS success.^16–18^ To account for this conformational heterogeneity directly, ensemble docking methods screen against multiple receptor structures from experimental or computational techniques to model target flexibility, which has been shown to improve early enrichment and hit diversity in GPCR VS campaigns.^18–22^

The quality of the ensemble structures used for screening, however, is critically dependent on the sampling method that was used to generate them. Ensemble structures generated from conventional MD (cMD) simulations face a fundamental timescale limitation in capturing allosteric conformational rearrangements, which occur on microsecond-to-second timescales.^23,24^ Enhanced sampling methods (e.g., metadynamics, accelerated MD, steered MD, random acceleration MD, etc.) improve the capture of GPCR allosteric dynamics by applying a biasing force or boost to overcome high energy barriers, which enables access to conformational states otherwise inaccessible to cMD.^25,26,27^ Gaussian accelerated MD (GaMD) is particularly attractive because these simulations may be run without prior knowledge of reaction coordinates and can yield thermodynamically re-weighted ensembles that faithfully reconstruct the free energy landscape of the allosteric pocket.^9,28,29^ We previously used GaMD-ensembles for retrospective docking of A1AR PAMs using AutoDock4, demonstrating that screening against these structural ensembles using the average binding energy (*BE_avg_*) of ligands across the ensemble for ranking improved docking performance over the cryo-EM structure.²⁸ However, the generalizability of this framework across multiple GPCR targets, diverse compound libraries (PAMs and NAMs), and docking programs, spanning empirical (Glide HTVS, AutoDock Vina), physics-based (DOCK3.8), and DL affinity prediction (Boltz-2) scoring functions, remains untested. It is also unknown whether Boltz-2 provides meaningful enrichment in this context if structural ensemble information can be meaningfully incorporated.

Here, we perform a systematic, multi-target retrospective benchmarking study of GPCR AM docking protocols. Using four model Class A GPCRs, the muscarinic M2 and M4 receptors (M2R, M4R), the β2-adrenergic receptor (β2AR), and C-C chemokine receptor type 2 (CCR2), we benchmark docking performance across three conventional docking programs (Schrödinger Glide, AutoDock Vina, and UCSF DOCK3.8), as well as Boltz-2, and assess PDB versus GaMD-derived ensemble performance in each context. We assess the impact of ensemble composition, size, and re-ranking strategy on early enrichment, as well as the impact of program-specific scoring function biases across protocols. The workflow, summarized in **Figure 1**, culminates in practical, dynamics-informed recommendations for GPCR AM docking protocol selection with the advent of high-accuracy enhanced sampling methods and DL-based prediction tools.

**Figure 1.**
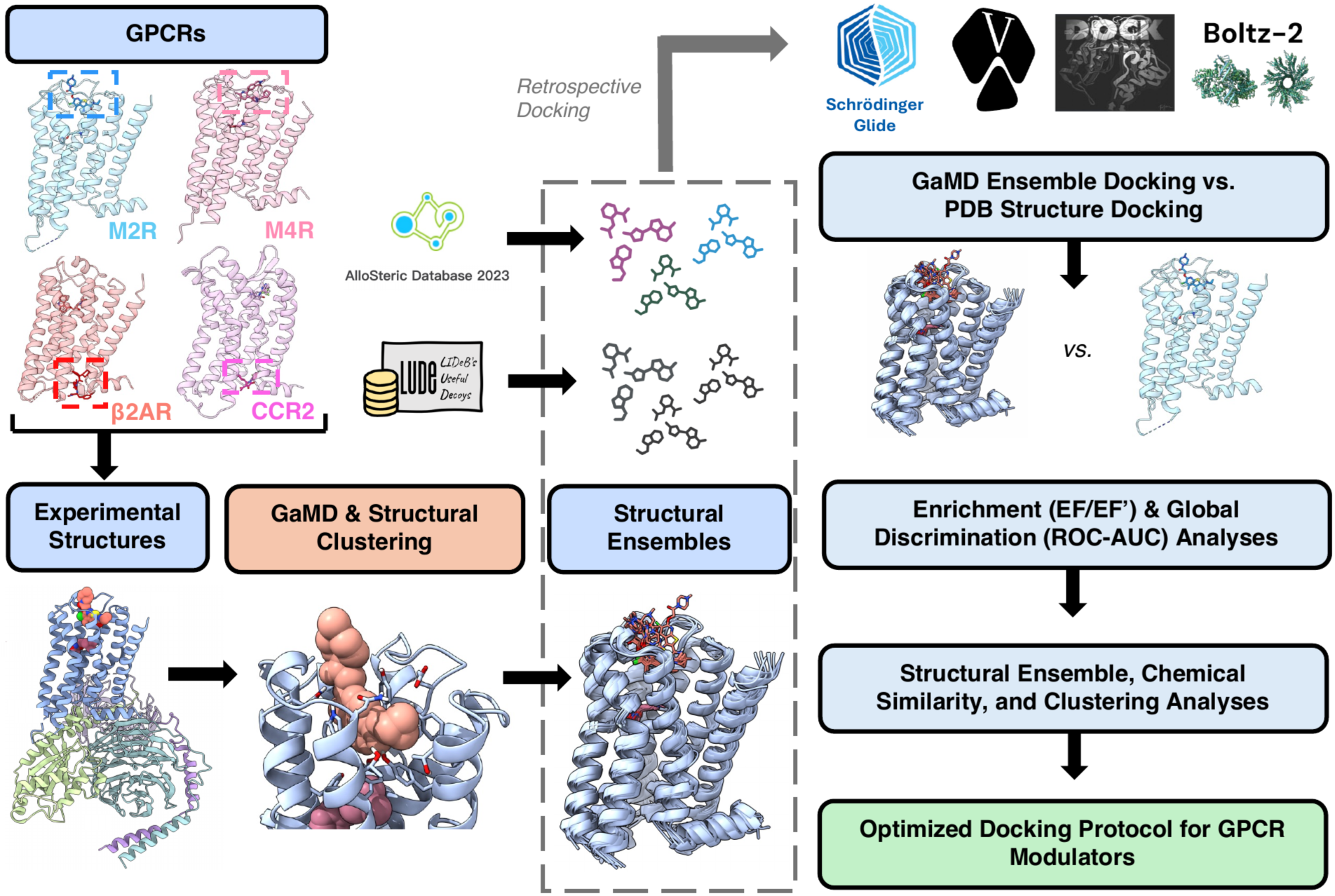
Workflow of Benchmarking Docking Protocols. Experimental (“PDB”) structures of four GPCRs (M2R, M4R, CCR2, β2AR) were used as starting points for Gaussian accelerated Molecular Dynamics (GaMD) simulations. GaMD frames were clustered for the allosteric pocket to generate receptor structural ensembles. Each ensemble was used for retrospective docking of known allosteric modulators against property-matched decoys using three popular docking programs, including Schrödinger Glide, AutoDock Vina, and DOCK3.8, as well as the DL co-folding program, Boltz-2. Docking scores or the score equivalent for Boltz-2 were aggregated across ensemble clusters using two re-ranking strategies (top or average) and benchmarked against PDB docking. Enrichment performance was evaluated using enrichment factors (EF), early EF (EF′), and ROC-AUC curves. Ensemble cluster structural variation was assessed using chemical descriptors (e.g., hydrophobicity, volume) and the Root Mean Square Deviation (RMSD). Chemical similarity and clustering of the top-ranking actives per program and docking method were also compared. The workflow culminates in an optimized docking protocol for the discovery of GPCR allosteric modulators.

## METHODS

### GaMD Theory and Energetic Reweighting

GaMD is an enhanced sampling method that uses a harmonic boost potential to flatten energy barriers and improve conformational sampling of biomolecules without predefined collective variables.^28^ Because this boost potential usually exhibits a near-Gaussian distribution, the canonical biomolecular free energy profiles can be accurately recovered through cumulant expansion to the second order. When the system potential *V*(*r⃑*) is lower than a threshold energy, *E*, a boost potential is added as

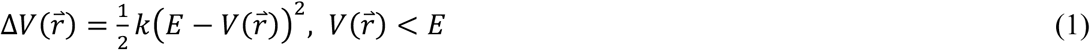

where *k* is the harmonic force constant. The modified system potential, *V*^∗^(*r⃑*) = *V*(*r⃑*) + Δ*V*(*r⃑*), is given by

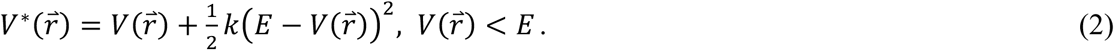

When the system is above the threshold energy, *V*(*r⃑*) ≥ *E*, the boost potential is set to zero and *V*^∗^(*r⃑*) = *V*(*r⃑*). The two adjustable simulation parameters, *E* and *k*, are determined based on three enhanced sampling criteria. The system threshold energy for applying the boost potential needs to be set within the following range:

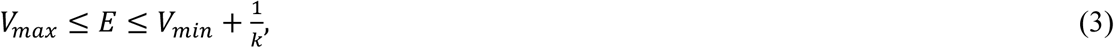

where *V*_*max*_ and *V*_*min*_ are the maximum and minimum potential energies of the system.^28^ In turn, for *k* to satisfy Eq. (3), it must meet the condition 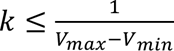. We define 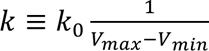 requiring 0 < *k*_0_ ≤ 1, where *k*_0_ sets the magnitude of the applied boost potential, i.e., a greater *k*_0_ value results in a higher applied boost potential to the system.

To ensure reliable energetic reweighting using cumulant expansion to the second order, the standard deviation of the boost potential Δ*V* must remain sufficiently small:

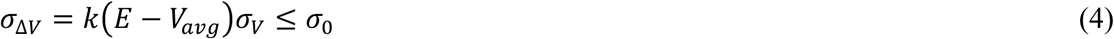

where *V*_*avg*_ and *σ*_*V*_ are the mean and standard deviation of the system potential energy, *σ*_Δ*V*_ is the standard deviation of Δ*V*, and *σ*_0_ is a user-defined upper limit (e.g., 10*k*_*B*_*T*).^30^ In this study, the threshold energy *E* was set to its lower bound, *E* = *V*_*max*_, and *E* and *k* were used to compute the magnitude of the applied boost potential:

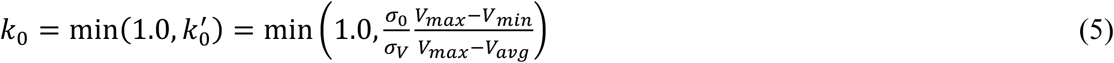

Here, the dual potential boost was used, including the total potential boost (Δ*V*_*P*_) and the dihedral potential boost (Δ*V*_*D*_), allowing for maximum simulation acceleration.^31^ As a result, the final simulation parameters consist of the threshold energy *E* and the force constants *k*_0*P*_ and *k*_0*D*_ for the total and dihedral potential boosts, respectively.

Per simulation, the potential of mean force (PMF) (“free energy”, in kcal/mol) was calculated considering the probability distribution along a reaction coordinate written as *p*^∗^(*A*).^32^ This can be rewritten to recover the canonical ensemble distribution, *p*(*A*), using the boost potential Δ*V*(*r⃑*) of each frame as:

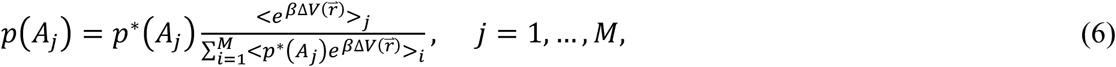

where *M* is the number of bins, *β* = *k*_*B*_*T*, and 〈*e*^*β*Δ*V*(*r⃑*)^〉_*j*_ is the ensemble-averaged Boltzmann factor of Δ*V*(*r⃑*) for frames in the *j*^*th*^ bin. Using cumulant expansion, the ensemble-averaged reweighting factor can be approximated:

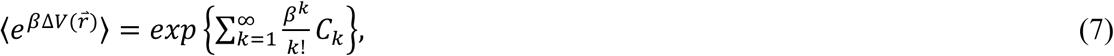

where the first two cumulants are

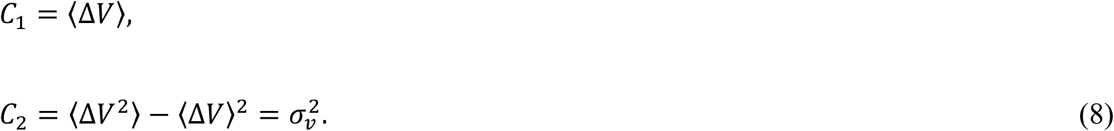

When the boost potential from GaMD simulations follows a near-Gaussian distribution, cumulant expansion to the second order provides a reliable approximation of the reweighting factor.^28,30^ Ultimately, the reweighted free energy *F*(*A*) = −*k*_*B*_*T* ln *p*(*A*) is calculated as:

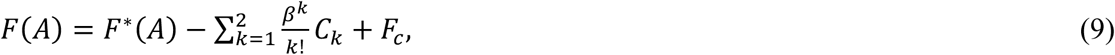

where *F*^∗^(*A*) = −*k*_*B*_*T* ln *p*^∗^(*A*) is the modified free energy extracted from the GaMD simulation and *F*_*c*_ is a constant.

### GaMD Simulations and Clustering for Structural Ensembles

GaMD simulations of the M2R, β2AR, and CCR2 were obtained from a previous study,^9^ and M4R from a more recent study using a similar protocol.^33^ A summary of the system setup and GaMD simulation protocol is provided here. Four experimental GPCR structures were used: the active-state Go-coupled M2R bound to the agonist Iperoxo and PAM LY2119620 (PDB: 6OIK),^34^ the active-state Go-coupled M4R bound to acetylcholine and PAM MK-97,^35^ the inactive-state β2AR bound to the antagonist Carazolol and NAM Cmpd-15PA (PDB: 5X7D),^36^ and the inactive-state CCR2 bound to the antagonist BMS-681 and NAM CCR2-RA-[R] (PDB: 5T1A).^37^ All water and heteroatom molecules except the ligands were removed from each structure, and chain termini were capped (ACE, N-terminus; CT3, C-terminus). Each complex was embedded in POPC lipid bilayers and solvated in 0.15 M NaCl. Finally, AMBER^38^ force field parameter sets were used: ff19SB^39^ for proteins, GAFF2 for ligands using the AM1-BCC charging method^40^, LIPID17 for lipids, and TIP3P^41^ for water. Periodic boundary conditions were applied to each simulation system. The SHAKE^42^ algorithm and a 2 fs time step were used; the temperature was kept at 310 K using the Langevin thermostat,^43,44^ and the pressure was kept constant at 1.0 bar using the Berendsen barostat.^45^ Electrostatic interactions were calculated using the particle mesh Ewald (PME) summation,^46^ and a long-range interactions cutoff of 9.0Å was used. Each system was minimized for 5000 steps with constant number, volume, and temperature (NVT) at 310 K, followed by further equilibration using constant number, pressure, and temperature (NPT) for 375 ps at 310 K. Next, 10-ns cMD simulations were performed with the NPT ensemble at 1 atm and 310 K to compute system potential statistics (*V*_*max*_, *V*_*min*_, *V*_*av*g_, and *σ*_*V*_). After 40-ns of GaMD equilibration with optimized acceleration parameters, three independent 500-ns dual-boost production runs with randomized velocities were performed per system using *E* = *V*_*max*_and an upper limit of 6.0 kcal/mol on the dihedral and total potential energetic terms.

Simulation frames from all independent trajectories (3×500ns) of a system were clustered using the hierarchical agglomerative algorithm implemented in CPPTRAJ.^47,48^ Clustering was performed using the heavy atoms of allosteric binding pocket residues (i.e., residues within 5Å from the PDB co-bound AM) with average linkage and a target ensemble of 10 clusters. With cumulant expansion to the second order, the per-frame boost potential was used to reweight the cluster populations to recover the original free energy of each cluster per system. Clusters with fewer than 500 frames and free energy above 8 kcal/mol were excluded as sparsely visited, high-noise states; only the M2R cluster 09 met this criterion, and all other clusters were retained for docking.

### Docking

#### Glide

Schrödinger Glide High-throughput Virtual Screening (HTVS) (version 2024-4) was used for PDB and structural ensemble docking.^49,50^ The default settings were used, with van der Waals radius scaling for the nonpolar ligand atoms set to 0.8 and a partial charge cutoff of 0.15. In addition, nonplanar amide conformations were penalized, the OPLS2005 force field was used, and five poses per ligand were subjected to post-docking minimization (top pose retained, ranked by GlideScore). Systems were prepared with the Schrödinger Protein Preparation Workflow (PPW):^51^ non-GPCR chains, non-ligand compounds, metals, and heteroatoms were removed, and side chains were filled. Hydrogens were added using Epik^52^ at a pH of 7.4 +/-2.0, hydrogen bond assignments were optimized using PROPKA,^53,54^ and restrained minimization was performed using the OPLS4 force field.^55^ Ligands were prepared in LigPrep (OPLS4) in which they were desalted, set with ionization states at pH 7.0 (Epik Classic), and tautomer/stereoisomer enumeration was performed (max 16 per ligand, max 500 atoms). The grid was centered on residues within 5 Å of the PDB co-bound modulator (inner box 10 Å; outer box 25 Å). The outer grid was extended to 36 Å for M4R to accommodate its largest ligand.

#### Vina

AutoDock Vina (“Vina”) (version 1.2.7) was used for benchmarking.^56,57^ Proteins were prepared using AutoDockTools (version 1.5.7), keeping only the GPCR receptor chain and orthosteric and allosteric ligands. All waters were deleted, polar hydrogen atoms were added, missing side-chain atoms were repaired, and Gasteiger charges were computed. Ligand SMILES were standardized for tautomers and protonation (pH 7.0, molscrub 0.1.1), enumerated and embedded with RDKit 2024.4.3, and converted to PDBQT with Meeko.^58^ The search space (30 Å) grid box center was defined around the PDB co-bound modulator. Docking used default parameters with an exhaustiveness of 8, generating 9 maximum binding poses per ligand, and an energy range of 3.0 kcal/mol.

#### DOCK3.8

UCSF DOCK3.8 (“DOCK3.8”) was used for docking as well, leveraging a physics-based score composed of three energy terms: van der Waals interaction, electrostatic interaction, and ligand desolvation on precomputed grids.^59,60^ Proteins were protonated in ChimeraX,^61^ and only the key components (i.e., GPCR and the allosteric and orthosteric ligands) were retained. DOCK3.8-compatible atom types and partial charges for the orthosteric ligand were added to the amb.crg.oxt and Prot.table.ambcrg.ambH parameterization files, respectively (as described in the DOCK3.7 wiki,^62^ and Bender et al.^60^). Matching spheres and scoring grids were generated with Blastermaster (pydock3)^63^: CHEMGRID (AMBER, vdW)^64^, QNIFFT (electrostatics)^65,66^, and SOLVMAP (desolvation)^67,68^. Matching spheres were generated using the PDB co-bound modulator heavy atoms converted to spheres via SPHGEN. These were supplemented by nearby receptor surface spheres until a total of 45 matching spheres was reached. Ligand .db2 files (DOCK3.8-compatible) were built from SMILES using build3d38 (TLDR); 10 M2R PAMs and 850 decoys failed to build.^63^ Because the M2R allosteric pocket is large and solvent-exposed, which makes it far harder to screen than a typical buried orthosteric site, the match goal was set to 5000, bump_maximum and bump_rigid were raised to 100.0, and internal clash checking was disabled to allow broader sampling. Up to 100 poses per ligand were scored, minimized, and written.

#### Boltz-2

The DL co-folding model, Boltz-2 (v2.2.1),^14^ was used for PDB and ensemble docking for its combined structural and affinity predictions. The default Boltz-2 settings were used, and input YAML configuration files were used to set the GPCR target sequence and compound SMILES string. Because Boltz-2 predicts from sequence rather than structure, GaMD ensemble clusters were supplied as structural templates, i.e., cleaned CIF files of the GPCR and co-bound ligand, with template forcing (force: true, threshold: 0.1 Å) to steer reverse diffusion so predicted backbones did not deviate from the template. Receptor sequences were set to the PDB reference, and the multiple-sequence alignments (MSA) were automatically computed for every protein chain using the mmseqs2 server.^69^ Boltz-2 has two main affinity outputs: (i) a binding likelihood classification-based value predicting if the provided ligand will bind to the target, set between 0 and 1, and (ii) a regression-based binding affinity prediction value in log10(IC50), based on an IC50 measured in μM. Previous work has benchmarked Boltz-2’s screening potential using each affinity output as an independent protein-ligand scoring and ranking metric made to be comparable to traditional exhaustive docking methods.^70,71^ Here, a combined scoring function (“CS”), as previously described in the original Boltz-2 manuscript, was used:^14^

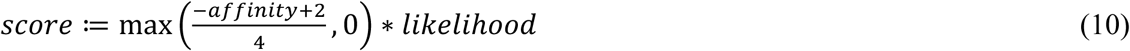

where the affinity prediction is approximately normalized, lower-bounded to zero, and weighted by the predicted binding likelihood. Only a single complex was generated per compound.

### Ensemble Scoring

The binding probability *p*_*i*_ of docking a ligand against a structural cluster *i* in the GaMD-derived receptor ensemble was weighted by the Boltzmann distribution of the clusters 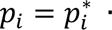 *e*^*PMF_i_*/*k_B_T*^, where *PMF*_*i*_ is the reweighted PMF of the ith structural cluster in the ensemble, *k*_*B*_ is the Boltzmann constant, and *T* is the temperature. From docking, the ligand binding probability 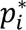 was related to the raw docking score *E*_*dock*_ as 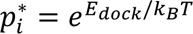. Given that *p*_*i*_ = *e*^*BE_i_*/*k_BT_*^ is defined by the ligand binding energy to the ith receptor structural cluster (*BE*_*i*_), the predicted binding energy for each receptor conformation in the ensemble was calculated as:

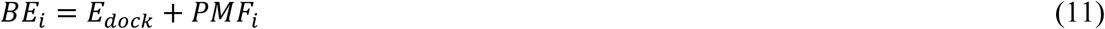

where *PMF*_*i*_ denotes the reweighted free energy associated with the ith structural cluster and *E*_*dock*_ represents the raw docking score. Compounds were ranked by the single best binding energy over all clusters,

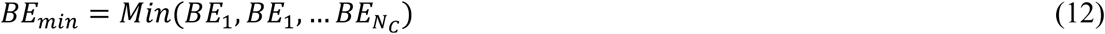

where *N*_*C*_ is the total number of clusters in the receptor ensemble, and the average across clusters:

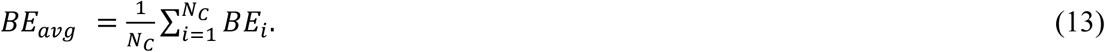

To extend the ensemble scoring framework to Boltz-2, which lacks a comparable binding-energy output, the CS (Eq. 10) was PMF-corrected. First, the Boltz-2 raw affinity prediction value *y*_*i*_ for a given compound was converted to *G_bind,i_* in kcal/mol using the Boltz-2 conversion formula^14^:

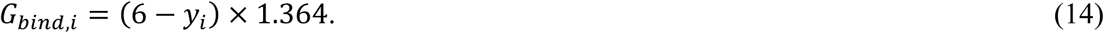

The PMF correction for the *i*th receptor structural cluster was then applied:

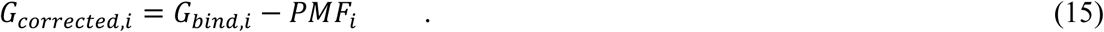

With the PMF-corrected free energy applied, this was expressed as a corrected affinity prediction in log10(IC50) units:

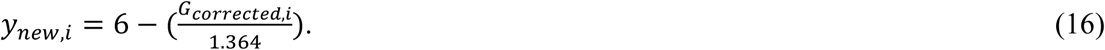

Unlike Eq. 11, where *PMF_i_* is added to a negative docking score, *PMF_i_* is subtracted from the positive *G_bind,i_* in Eq. 15. Boltz-2 yields a positive quantity, such that subtraction equivalently results in a decrease of *G_corrected,i_* and thus an increase of *y_new,i_* via Eq. 16, producing a lower CS for high-energy receptor conformations and preserving the intended thermodynamic reweighting values of high-PMF clusters. The corrected score is:

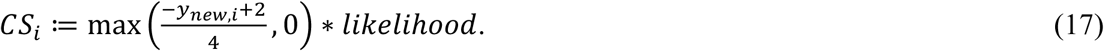

Boltz-2 ensemble predictions were ranked by the maximum (*CS*_*max*_) and average (*CS*_*av*g_) score across clusters, analogous to *BE*_*min*_and *BE*_*av*g_.

### Docking Enrichment and Validation

The docking performance of each protocol was assessed by its ability to rank true binders, i.e., AMs, higher than decoys, using enrichment factors (EF) at a given sampled percentile (X%) with an emphasis on the earliest percentiles (0.5-5%):

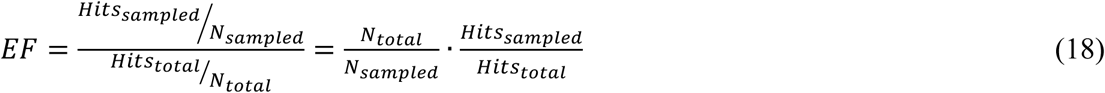

where *N*_*total*_ is the number of total compounds in the GPCR target library, *N*_*sampled*_ is the number of compounds used for ranking at a given percentile sampled (X%), *Hits*_*total*_ is the number of total hits, i.e., AMs, in the GPCR target library, and *Hits*_*sampled*_ is the number of hits found among the ranked compounds at a given percentile sampled. The early EF (EF’), which emphasizes the relative rank positions of *Hits*_*sampled*_, is calculated by:

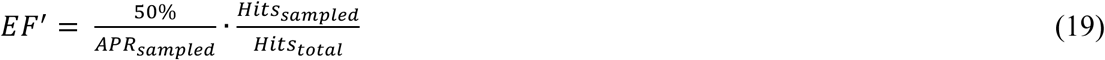

where *APR*_*sampled*_is the average percentile rank of the *Hits*_*sampled*_. In addition to EF and EF’, receiver-operator characteristic (ROC) curves and the area under the curve (AUC) values were used to measure overall protocol binder-decoy discrimination capabilities. Semilogarithmic ROC curves and the logAUC values were also calculated. Here, logAUC is the area under the semi-logarithmic ROC curve normalized to a random baseline, with higher logAUC values indicating greater early enrichment.^60^

### Compound Libraries

Compound libraries were prepared for each GPCR target by collecting known AMs from ASD2023; NAMs and PAMs are classified as inhibitors and activators, respectively.^11,72^ Each target library was manually curated, and inappropriate entries were removed. For β2AR, alprenolol, the well-known β2AR antagonist, was excluded from the library. For CCR2, a small peptide (LGTFLKC) was removed. For M2R, an M1-selective PAM was excluded, and for M4R, seven entries with missing records and one lacking a 3D structure were removed. SMILES were then standardized in RDKit 2025.3.6 using the following steps: explicit-hydrogen removal, metal disconnection, normalization/reionization, largest-fragment extraction, charge neutralization, and tautomer canonicalization. Duplicates were removed by standardized SMILES and charge-stripped InChI. Next, property-matched decoys were built at 50 per active. Three decoy generation protocols were initially tested: DUD-E,^73^ DUDE-Z,^74^, and LUDe.^75^ LUDe was chosen as it was the only protocol to consistently reach 50:1 across all targets and run both rapidly and locally. **Table 1** summarizes the per-target active and decoy counts and their chemical descriptors, as well as the median decoy-to-active similarity values using an ECFP4-based (2048 bits) Tanimoto similarity coefficient (Tc) metric, computed using RDKit 2025.03.6.

**Table 1.** Compound Libraries and Their Descriptors.^a^. ^a^We report the median values for all chemical descriptors and similarity metrics. Decoy-to-active TC (Tanimoto coefficient) is the median ECFP4-based (2048 bits) similarity between a target’s decoy library and active library.

| Target | Label | Total Count | MW | logP | HBD | HBA | RB | Charge | Rings | Decoy-to-active TC |
| --- | --- | --- | --- | --- | --- | --- | --- | --- | --- | --- |
| *M2R* | Actives | 230 | 548.728 | 4.162 | 0 | 4 | 7 | 1 | 5 | 0.093 |
|  | Decoys | 11,550 | 521.384 | 3.841 | 1 | 6 | 7 | 0 | 4 |  |
| *M4R* | Actives | 2314 | 385.684 | 3.443 | 1 | 6 | 4 | 0 | 4 | 0.100 |
|  | Decoys | 115,643 | 380.425 | 3.389 | 1 | 5 | 4 | 0 | 3 |  |
| *β2AR* | Actives | 64 | 388.576 | 5.396 | 2 | 4 | 4 | 0 | 4 | 0.095 |
|  | Decoys | 3200 | 390.428 | 4.991 | 1 | 4 | 5 | 0 | 3 |  |
| *CCR2* | Actives | 16 | 469.136 | 4.282 | 1.5 | 6 | 7.5 | 0 | 4 | 0.108 |
|  | Decoys | 800 | 462.503 | 3.878 | 1 | 6 | 6 | 0 | 4 |  |

### Structural and Chemical Analyses

GaMD-derived structural ensembles were inspected using CPPTRAJ clustering summaries: (i) pocket root-mean-square deviation (RMSD) over time, (ii) cluster assignment over time, (iii) frame count per cluster, (iv) fraction of total frames, (v) average intra-cluster RMSD, and (vi) average distance to centroid.^47,48^ Binding pocket properties for PDB structures and ensemble clusters were computed using Schrödinger SiteMap^76,77^ with the default parameters (a minimum of 15 site points in the initial site-finding stage, a more restrictive definition of hydrophobicity, a standard 0.7 Å grid, and cropped sitemaps at 4 Å) to evaluate the pocket surrounding the bound AM. In addition, the RMSD of the allosteric pocket in GaMD clusters relative to the first cluster was computed. Physicochemical descriptors (MW, logP, rotatable bonds, HBA, HBD, ring count, formal charge, heavy-atom count) of the compound libraries were assessed using RDKit 2025.3.6. In addition, the topological similarity to the PDB-bound modulator was assessed using Morgan fingerprints (radius = 2, 2048 bits) and pairwise Tc. Finally, BitBIRCH^78^ clustering of the union of top-5% M2R actives across protocols (RDKit circular fingerprints, Tanimoto threshold 0.65) yielded 14 clusters (F1-F14), and ECFP4 fingerprints of all M2R actives and decoys were projected onto two principal components fit on the combined matrix. The average Tanimoto similarity (iSIM) was used to assess intra-cluster scaffold similarity.

## RESULTS

### GaMD Simulations Yield Diverse GPCR Structural Ensembles

Initial benchmarking was performed using the cryo-EM structure and simulation clusters of the Go-coupled M2R bound to the agonist Iperoxo and PAM LY2119620 (PDB: 6OIK) (**Fig. 2a**).^34^ M2R PAMs bind to a solvent-exposed, shallow pocket located in the extracellular vestibule of the receptor, distinct from the buried orthosteric pocket, providing a challenging screening target for any conventional docking program. Nine representative structures of the M2R allosteric pocket were obtained from RMSD-based clustering from each independent all-atom, dual-boost GaMD simulation (3×500ns) of the M2R PDB (**Fig. 2b**).^9^ Over the course of the simulation, the allosteric pocket side-chain RMSD relative to the M2R PDB reference fluctuated between 2.1 ± 0.3 Å, indicating meaningful conformational sampling of the allosteric site (**Fig. S1a**). Clusters 00-03 were the most frequently visited states and were sampled throughout the entirety of the simulations (**Fig. S1b**). Overall, the per-cluster frame populations ranged from 35% (cluster 00) to just under 1% (cluster 08), while later clusters (cluster 04 and beyond) represented sparsely sampled, increasingly rare conformations (**Fig. 2c**). The resulting GaMD-reweighted free energy (kcal/mol) of each cluster indicated energetically favorable states for all clusters despite dramatic population differences, with a steep increase in free energy from cluster 04. These free energies allow us to reweight each compound’s raw docking score by the receptor’s probability of occupying each state, yielding a re-scoring scheme that reflects modulator binding more realistically than a single PDB structure.

**Figure 2.**
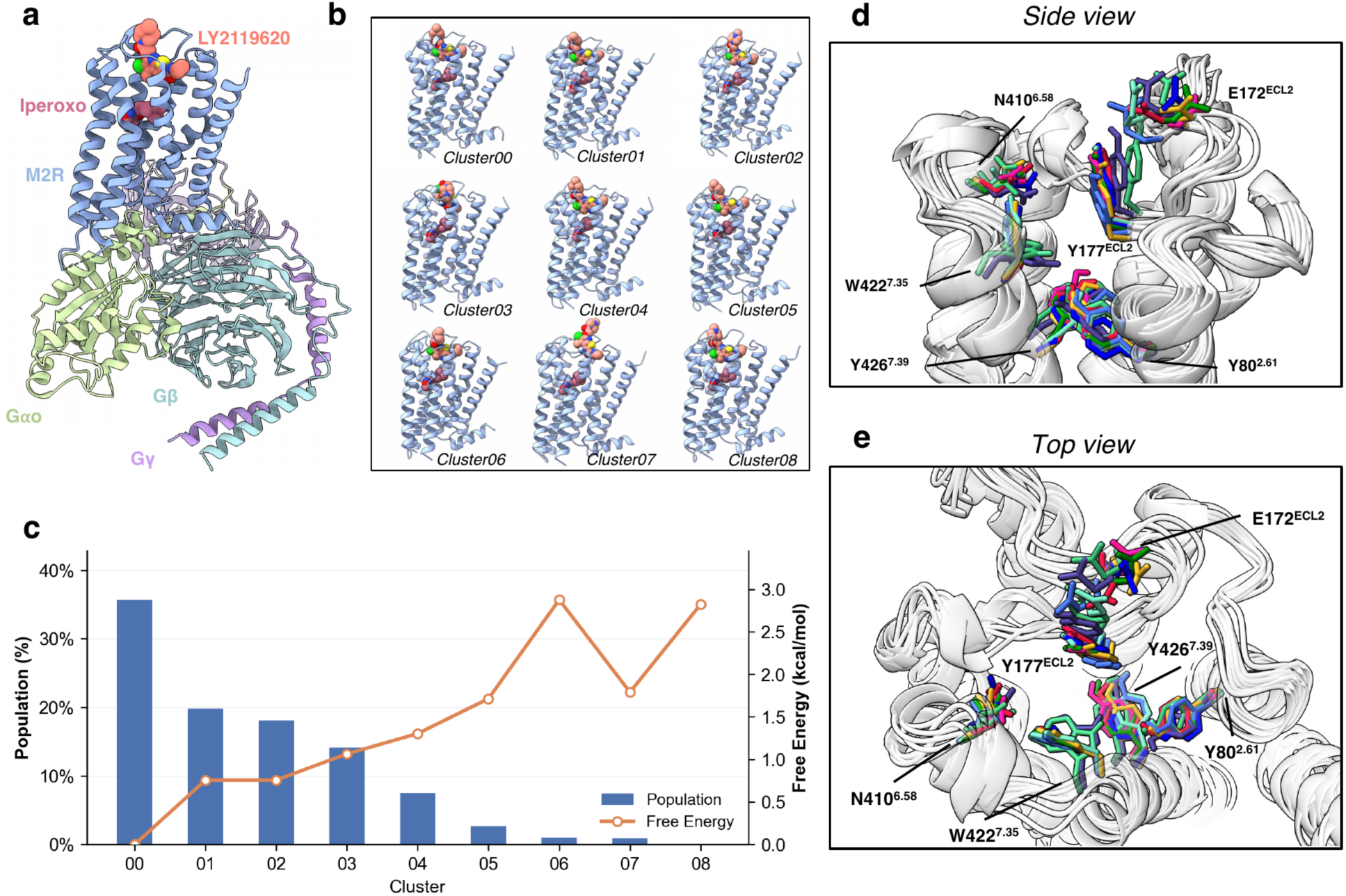
GaMD Structural Ensemble and Allosteric Pocket Diversity of the M2R Model System (a) Overview of the M2R-Go-iperoxo-LY2119620 complex structure. (b) Nine top-ranked structural clusters. (c) Cluster population (%, blue bars; left axis) and the reweighted free energy (kcal/mol, orange line; right axis). (d, e) Zoom-in side and top views of the M2R allosteric pocket highlight conformations of key residues: E172^ECL2^, Y177^ECL2^, Y80^2.61^, N410^6.58^, W422^7.35^, and Y426^7.39^.

Superposition of the representative frames from each of the nine clusters reveals considerable side-chain and backbone variation (**Fig. 2d-e**). The M2R allosteric pocket is largely preformed before modulator binding and further contracts when the modulator is bound, which is well-replicated in our ensembles. Notably, in multiple frames, W422^7.35^ adopts a horizontal rotamer position rather than the typical vertical conformation it takes when forming aromatic stacking with LY2119620. There were also considerable shifts in key loop regions, E172^ECL2^ and Y177^ECL2^, consistent with the known flexibility of ECL2 and its stabilizing interactions with LY2119620 (aromatic stacking, charge-charge piperidine interactions). SiteMap analysis showed the pocket volume varied substantially across the ensemble: from 94 Å^3^ (cluster 05) to 362 Å^3^ (cluster 06) (**Table 2**). Two clusters deviated markedly from both the PDB and the ensemble: cluster 05, with a small, extremely hydrophobic pocket (hydrophobic 2.556, balance 3.663, 93.98 Å^3^), and cluster 08, scored as a non-binding site (SiteScore 0.729, <0.8 threshold; 170 Å^3^). These ensembles successfully capture conformational and druggability variation across the M2R allosteric pocket that a PDB structure alone cannot capture.

**Table 2.** SiteMap Binding Pocket Properties for the M2R PDB and Structural Clusters.^a^. ^a^ *SiteScore* = quantifies molecule-binding vs. non-molecule-binding sites (>1.0 is a promising binder; 1.0 > SiteScore > 0.8 is a binding site; < 0.8 is a non-binding site), *DScore* = druggability score (>1.0 is a likely drug binder), *Balance* = the ratio of philic/phobic regions in the site (>0.5 considered good), *Hydrophobic/Hydrophilic* = hydrophobic/hydrophilic character of the site (average score for tight binding site is 1.0), *Don/Acc* = degree of hydrogen bond donor/acceptor character.^76,77^

| M2R Structure | SiteScore | DScore | Balance | Volume [Å<sup>3</sup>] | Hydrophobic | Hydrophilic | Don/Acc |
| --- | --- | --- | --- | --- | --- | --- | --- |
| Cluster00 | 1.084 | 1.113 | 1.481 | 290.18 | 1.369 | 0.925 | 0.812 |
| Cluster01 | 1.071 | 1.119 | 1.480 | 272.00 | 1.200 | 1.200 | 0.541 |
| Cluster02 | 1.112 | 1.156 | 1.889 | 315.56 | 1.527 | 0.808 | 0.715 |
| Cluster03 | 1.080 | 1.133 | 1.172 | 320.70 | 0.903 | 0.771 | 0.830 |
| Cluster04 | 1.099 | 1.099 | 3.663 | 93.98 | 2.556 | 0.698 | 0.513 |
| Cluster05 | 1.094 | 1.124 | 1.410 | 362.21 | 1.280 | 0.908 | 1.410 |
| Cluster06 | 1.060 | 1.087 | 1.253 | 237.36 | 1.191 | 0.951 | 0.542 |
| Cluster07 | 0.729 | 0.684 | 0.538 | 170.13 | 0.462 | 0.858 | 0.705 |
| Cluster08 | 1.070 | 1.119 | 1.423 | 330.65 | 1.134 | 0.797 | 0.698 |
| PDB | 1.068 | 1.107 | 1.139 | 322.08 | 0.985 | 0.865 | 0.893 |

### M2R Structural Ensembles Improve Enrichment and Rank-placement of Actives

Three conventional docking programs (Glide, Vina, DOCK3.8) and Boltz-2 were used to evaluate whether GaMD-derived conformational ensembles improved early AM enrichment relative to single PDB structures. Known M2R PAMs (n=230) and property-matched decoys (n=11,500) were docked across all M2R structures, and ensemble performance was assessed using either the top or average reweighted binding score across clusters. GaMD ensemble docking broadly outperformed PDB docking, with 18 of 24 early enrichment metrics (EF, EF′, AUC, and logAUC at 0.5-1%; 6 per program) improving across all four programs (**Table 3**). The conformational diversity of the ensemble yielded meaningful screening benefits for the majority of the protocols evaluated. The 5 (20.83%) metrics that did not improve were confined entirely to Boltz-2, where PDB and ensemble performance were already comparable (PDB AUC: 63.47%; *CS_max_* AUC: 63.32%), suggesting Boltz-2 affinity predictions were largely insensitive to the structural templates provided. Most notably, ensemble gains were most pronounced at the earliest percentiles, 0.5% and 1%, where nearly all (93.75%) EF and EF’ metrics improved over the PDB. Because only top-ranked hits (0.01-0.5%) in a prospective screen would typically advance to experimental follow-up, this early enrichment improvement provided by the structural ensemble is practically meaningful. Between all programs, Glide *BE_avg_* yielded the best EF’ across all sampled percentiles (**Table S1**), reflecting both strong early enrichment and high rank placement of M2R actives. The ensemble models appeared more robust in improving EF’ performance regardless of program. We found that the primary benefit of M2R ensemble docking was in improving the relative rank placement of actives rather than just increasing the number recovered at a given cutoff.

**Table 3.** Enrichment Factors and AUC Values from PDB and Ensemble Docking of M2R Allosteric Modulators Using Glide, Vina, DOCK3.8, and Boltz-2.^a^. ^a^EF = enrichment factor, EF’ = early enrichment factor, *BE_min_* = minimum binding energy, *BE_avg_* = average binding energy, *CS_max_* = maximum combined score, *CS_avg_* = average combined score. The best metric (EF/EF’/AUC/logAUC) within each program and between the targets is in bold.

| Program | Target | EF(0.5%) | EF'(0.5%) | EF(1%) | EF'(1%) | %AUC | %logAUC |
| --- | --- | --- | --- | --- | --- | --- | --- |
| *Glide HTVS* | PDB | 1.998 | 2.938 | 2.996 | 2.378 | 59.22 | 5.95 |
| | Ensemble ($BE_{min}$) | 11.985 | 10.101 | **10.792** | 10.870 | 56.63 | 8.62 |
| | Ensemble ($BE_{avg}$) | **18.833** | **20.136** | **13.382** | **18.261** | 56.43 | **9.93** |
| *AutoDock Vina* | PDB | 12.030 | 10.218 | 10.312 | 11.548 | 46.77 | 4.74 |
| | Ensemble ($BE_{min}$) | **16.327** | **16.629** | **12.030** | **15.555** | 49.81 | **7.57** |
| | Ensemble ($BE_{avg}$) | 6.874 | 5.128 | 6.874 | 5.901 | 49.31 | 4.23 |
| *DOCK3.8* | PDB | 1.913 | 2.623 | 2.416 | 2.362 | 61.92 | **6.07** |
| | Ensemble ($BE_{min}$) | **4.746** | **4.250** | 2.877 | **3.694** | 59.09 | 5.41 |
| | Ensemble ($BE_{avg}$) | 3.796 | 3.169 | **3.357** | 3.543 | 54.03 | 4.02 |
| *Boltz-2* | PDB | 3.423 | 2.831 | **5.614** | 4.337 | 63.47 | **11.29** |
| | Ensemble ($CS_{max}$) | 4.317 | 4.162 | 3.886 | 3.738 | 63.32 | 10.71 |
| | Ensemble ($CS_{avg}$) | **5.181** | **4.878** | 5.181 | **4.878** | 62.79 | 10.90 |

PDB docking performance varied substantially across programs: Vina outperformed Glide, Boltz-2, and DOCK3.8, by up to 6-fold at 0.5% sampled (**Fig. 3a**). Vina’s strong PDB performance outperformed all early percentile docking metrics from both PDB and ensemble models of DOCK3.8 and Boltz-2. The absolute enrichment values were therefore partly determined by program-specific scoring function handling of the M2R PAMs, independent of ensemble benefit. Only Glide (*BE_avg_*) and Vina (*BE_min_*) ensembles surpassed Vina’s PDB performance. Program performance rank-order inverted when assessed by logAUC (**Fig. 4**): Boltz-2 (11.29%) and DOCK3.8 (6.07%) overtook both Glide (5.95%) and Vina (4.74%). Boltz-2 and DOCK3.8 accumulated enrichment more gradually across early percentiles rather than concentrating it at the earliest ranks. Practically, however, EF and EF′ at fixed cutoffs (0.5-1%) more directly reflect hit prioritization utility than logAUC, which integrates performance across percentiles. Therefore, logAUC ranking can obscure poor performance at the earliest of percentiles, as demonstrated by Boltz-2 and DOCK3.8.

**Figure 3.**
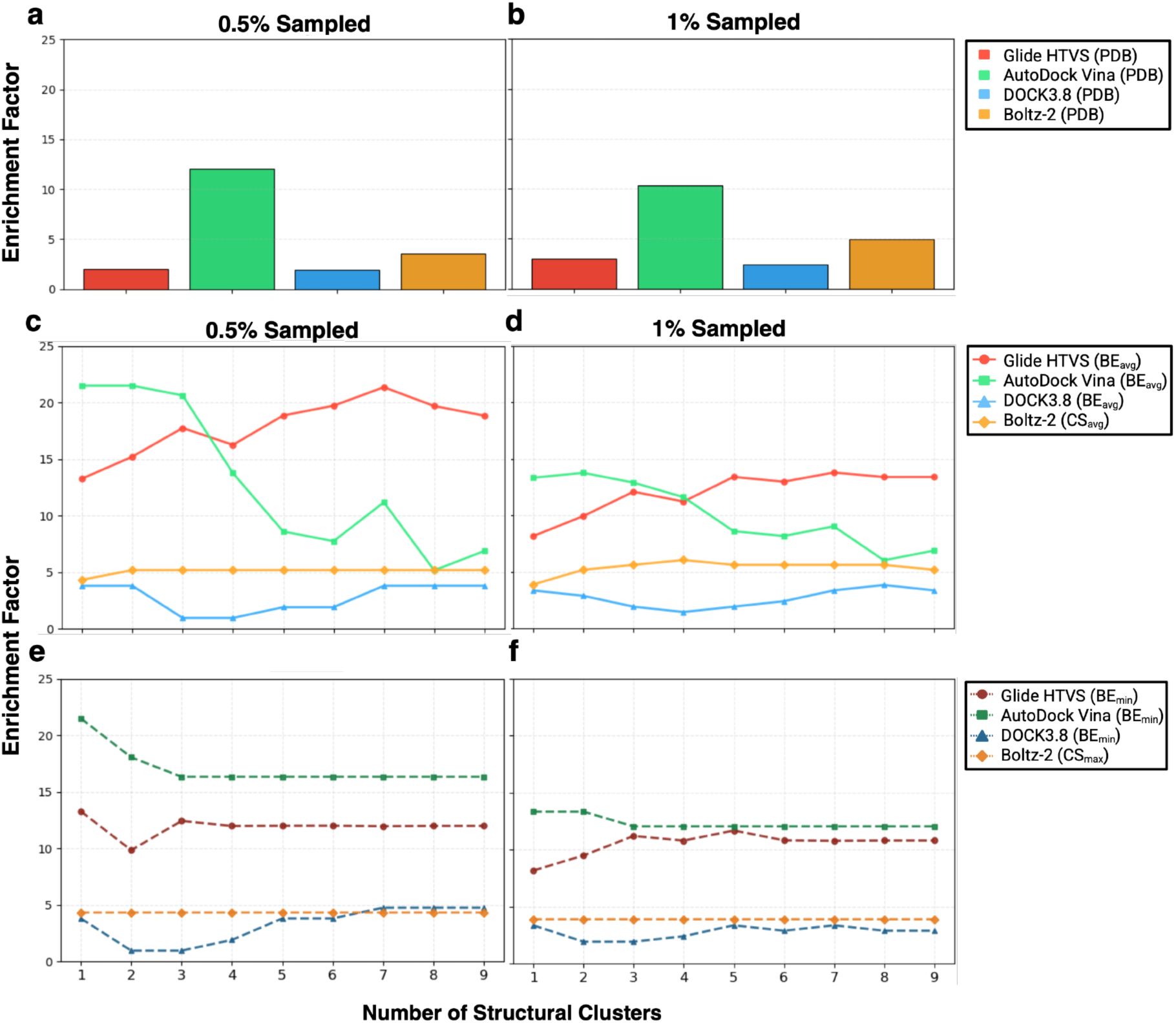
Performance Comparison of Different Docking Programs at M2R. (a-b) Docking Enrichment Factors (EF) from the top (a) 0.5% and (b) 1% of ranked ligands using the M2R PDB structure. (c-d) Ensemble EFs, by program, using the average ensemble score re-ranking method (*BE_avg_* [Glide, DOCK, Vina], *CS_avg_* [Boltz-2]). (e-f) Ensemble EFs, by program, using the best ensemble score re-ranking method (*BE_min_* [Glide, DOCK, Vina], *CS_max_* [Boltz-2]). Ensemble enrichment plots are plotted as a function of ensemble size (# of structural clusters included for EF calculations).

**Figure 4.**
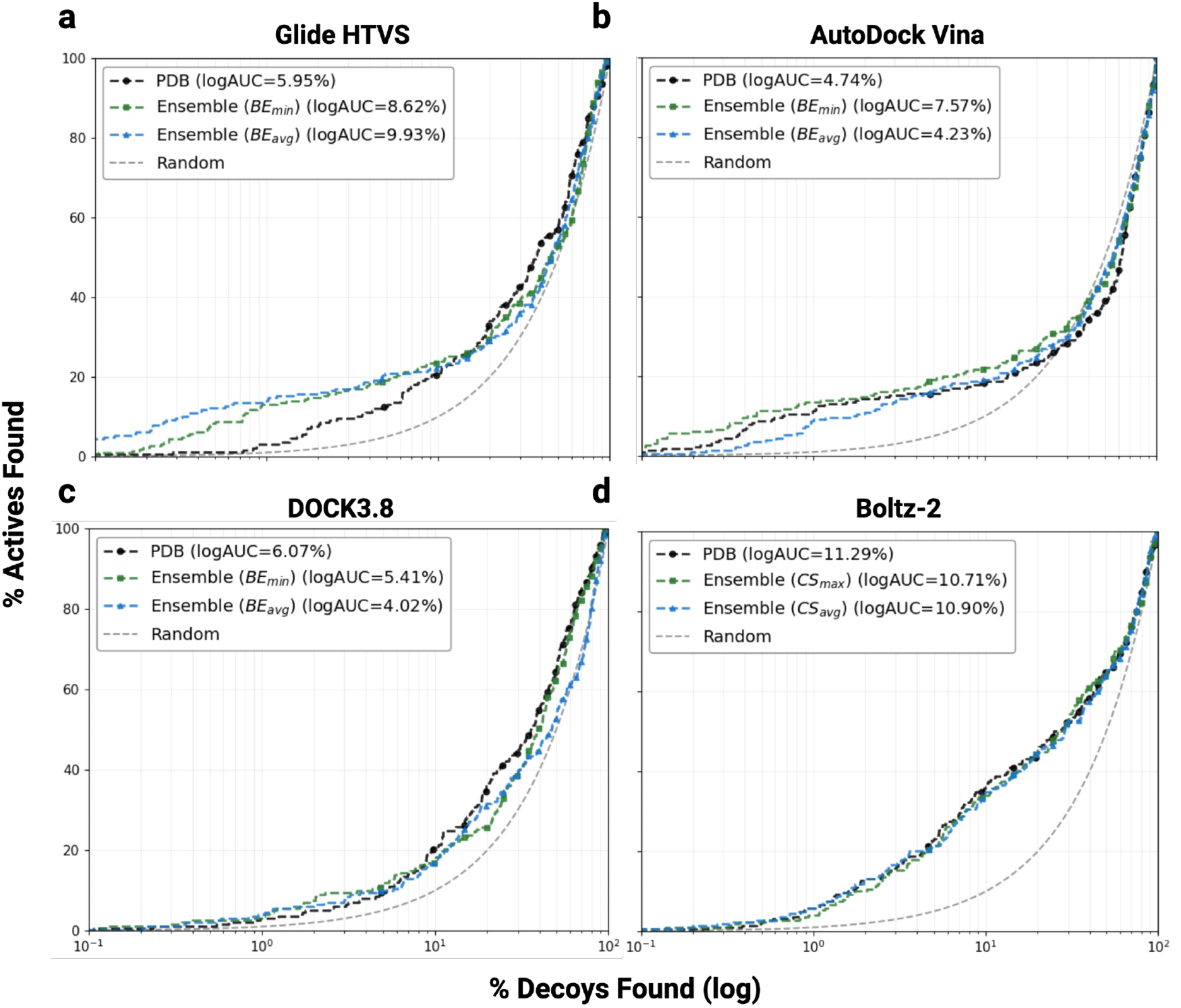
ROC Curves of Allosteric Modulator Docking at M2R. (a-d) Semilogarithmic ROC curves and the (log) Area Under the Curve (AUC) values for the PDB (solid black) and ensemble docking (*BE_min_* and *CS_max_* [green, top ensemble docking score]; *BE_avg_* and *CS_avg_* [blue, average ensemble docking score]) by program: Glide, Vina, DOCK3.8, and Boltz-2. The x-axis is plotted on a log scale to emphasize early enrichment performance. logAUC is the area under the semi-logarithmic ROC curve normalized to the random baseline (grey dashed line); higher logAUC indicates greater early enrichment.

Despite Vina’s superior early enrichment, its AUC (49.81%) approached random discrimination. Boltz-2, however, yielded the highest AUC values in different settings (PDB: 63.47%; Ensemble with *CS_max_*: 63.32%) and competitive late-percentile EFs (3-5%) (**Tables 3, S1)**. This indicates that Boltz-2 captured broad active-decoy separation but failed to concentrate actives at the earliest ranks, matching its poor ensemble EF and EF’ performance. Ensemble inclusion benefited Glide and Vina most substantially, wherein all 12 EF and EF′ metrics favored Glide *BE_avg_* and 11 of 12 favored Vina *BE_min_*, improving by up to ∼15 enrichment points relative to the PDB. Glide yielded the best overall M2R ensemble EF and EF′, closely followed by Vina, then Boltz-2 and DOCK3.8. Glide demonstrated the most consistent improvement over its own PDB baseline and the strongest early rank placement of actives across re-ranking strategies, making itself and Vina the two most reliable ensemble docking programs in this M2R PAM screening.

### Optimal Ensemble Re-ranking Strategy Is Program-Dependent and Influenced by GaMD Reweighted Free Energy Values

To determine how the number of M2R structural clusters and their GaMD-reweighted free energies jointly influence early PAM enrichment, we systematically varied the number of clusters included in ensemble docking and evaluated both *BE_min_* and *BE_avg_* re-ranking strategies across all four programs (**Fig. 3c-f**). For Glide *BE_avg_*, enrichment increased steadily with each additional cluster, particularly clusters 00-02. These clusters collectively account for ∼87% of sampled frames and span a compressed free energy range of only 0-2.83 kcal/mol (**Fig. 2c**). Averaging across these thermodynamically similar, well-populated conformations yielded a meaningful performance improvement. The opposite trend was observed for Vina *BE_avg_*, where a dramatic enrichment decline occurred upon introduction of clusters 03 and 04, wherein steeper free energy penalties from GaMD reweighting began to outweigh Vina’s docking scores for those conformations and resulted in a diluted enrichment signal. Glide’s scoring function appeared to be more robust than Vina to these free energy differences. When *BE_min_* was used across Vina and Glide, EF peaked at cluster 00 and gradually plateaued by cluster 03 with 0.5% sampled (**Fig. 3e**). The dominant ground-state conformation drove the best per-compound scores across the ensemble, and additional clusters contributed diminishing marginal benefit. DOCK3.8 and Boltz-2 were substantially less sensitive to ensemble size.

DOCK3.8 *BE_min_* and *BE_avg_* benefited modestly from the inclusion of clusters 04-08, while Boltz-2 enrichment remained entirely flat across all cluster numbers and percentiles (**Fig. 3c-f**). This insensitivity likely reflects convergence of Boltz-2’s reverse diffusion model toward the cryo-EM ground state regardless of the cluster template provided. Such results indicated that the conformational diversity of the M2R ensemble was marginally productive when assessed by DOCK3.8’s scoring and practically irrelevant to Boltz-2’s affinity predictions. We find that the optimal ensemble re-ranking strategy is entirely program-dependent, e.g., *BE_avg_* for Glide and *BE_min_* for Vina, and handling and scoring of the structural clusters cannot be assumed transferable across programs without prior retrospective evaluation.

### Program and Ensemble Re-ranking Strategies Determine Chemical Diversity of Recovered Actives

Beyond enrichment performance, the choice of program and ensemble re-ranking strategy appeared to determine which chemically distinct M2R PAM scaffolds were recovered. We first examined whether *BE_min_*-alone, *BE_avg_*-alone, or shared ensemble re-ranking yielded differences in the total percentage of actives recovered at the earliest percentiles. At 0.5% sampled, *BE_avg_*-alone recovered the highest percentage of actives for Glide, while *BE_min_*-alone dominated for Vina and DOCK3.8, and *CS_max_*-alone or *CS_avg_*-alone performed equivalently for Boltz-2 (**Fig. 5**). This is consistent with the program-dependent re-ranking behavior established in the previous section. For DOCK3.8 and Boltz-2, both re-ranking methods produced comparable and substantially lower enrichment than Vina and Glide, being largely insensitive to re-ranking strategy. Critically, by 3% sampled, nearly all actives recovered by either method alone were shared, regardless of program, motivating a consensus approach for ensemble prospective screening applications. Taking the union of hits ranked highly by both *BE_min_* and *BE_avg_* maximizes the chemical diversity of the earliest-percentile hit pool without discarding actives uniquely identified by either strategy alone.

**Figure 5.**
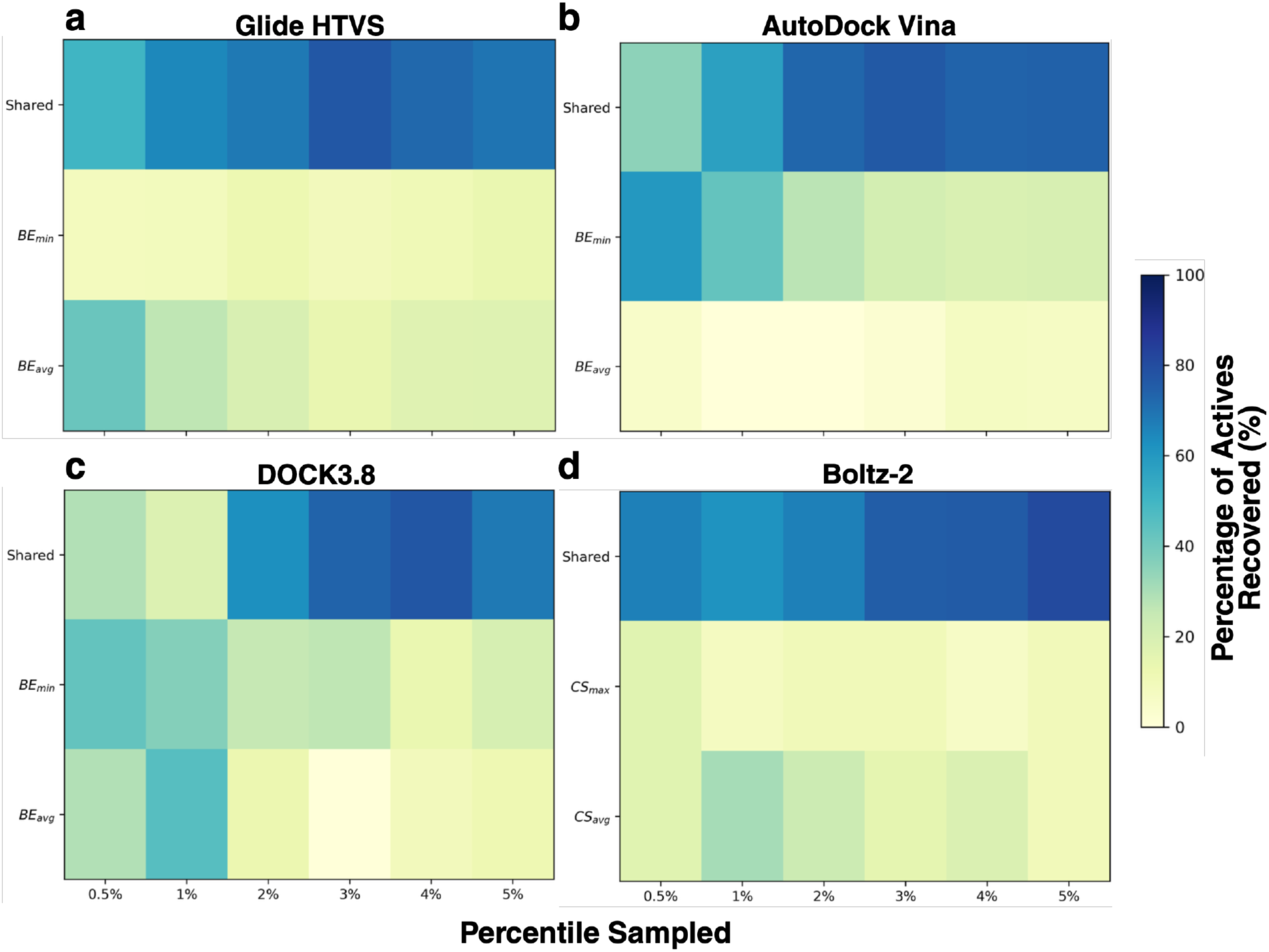
Percentage of M2R Actives Recovered by Top, Average, or Shared Re-ranking Methods Across Early Sampled Percentiles. (a-d) The percentage of actives recovered at each sampled percentile (0.5-5%) using the average (*BE_avg_* [Glide, DOCK, Vina], *CS_avg_* [Boltz-2]) or top (*BE_min_* [Glide, DOCK, Vina], *CS_max_* [Boltz-2]) ensemble docking scores for ranking, compared with the shared overlap (i.e., actives enriched by both ensemble scoring methods). M2R ensemble docking active recovery percentages at 0.5-5% sampled from (a) Glide HTVS, (b) AutoDock Vina, (c) DOCK3.8, and (d) Boltz-2.

Next, to characterize the scaffold diversity of recovered actives, BitBIRCH clustering was performed on the union of M2R actives recovered across all programs and protocols at 0.5-5% sampled (n=116), yielding 14 distinct chemical families (F1-F14) (**Fig. 6, S6**). Three families were identified in total; four were identified exclusively by DOCK3.8 and not by other programs, and one was exclusively identified by Boltz-2. The program selection used for M2R PAM screening, therefore, determined what regions of chemical space were accessible and enriched at the top percentiles. The two predominant families, F1 and F2 (**Fig. 6e-f**), exhibited distinct program-specific recovery patterns: Glide and Vina preferentially recovered the more structurally similar F2 scaffolds (iSIM 0.853), while DOCK3.8 and Boltz-2 preferentially recovered the more diverse F1 scaffolds (iSIM 0.658), and DOCK3.8 recovered no F2 actives at these top percentiles (**Fig. 6a-d**). Surprisingly, Boltz-2 uniquely captured the broadest scaffold distribution, recovering actives from the four largest families with near-equal representation across PDB and ensemble docking protocols. Boltz-2 likely prioritizes different molecular features than empirical or physics-based scoring functions, recovering chemically diverse actives that conventional programs deprioritize, albeit at lower overall enrichment. Beyond scaffold composition, per-method hit counts varied substantially: Boltz-2 with *CS_avg_* identified the most hits (n=55), while DOCK3.8 identified the fewest regardless of method (PDB: 18; *BE_min_*: 22; *BE_avg_*: 20) (Fig. **6c-d**). Taken together, while the majority of actives recovered within a program converged at later percentiles, each program preferentially enriched chemically distinct scaffolds at the earliest percentiles (0.5-1%), and GaMD ensemble docking recovered actives across scaffold families that PDB-only screening would have missed entirely (**Fig. 6e-g**).

**Figure 6.**
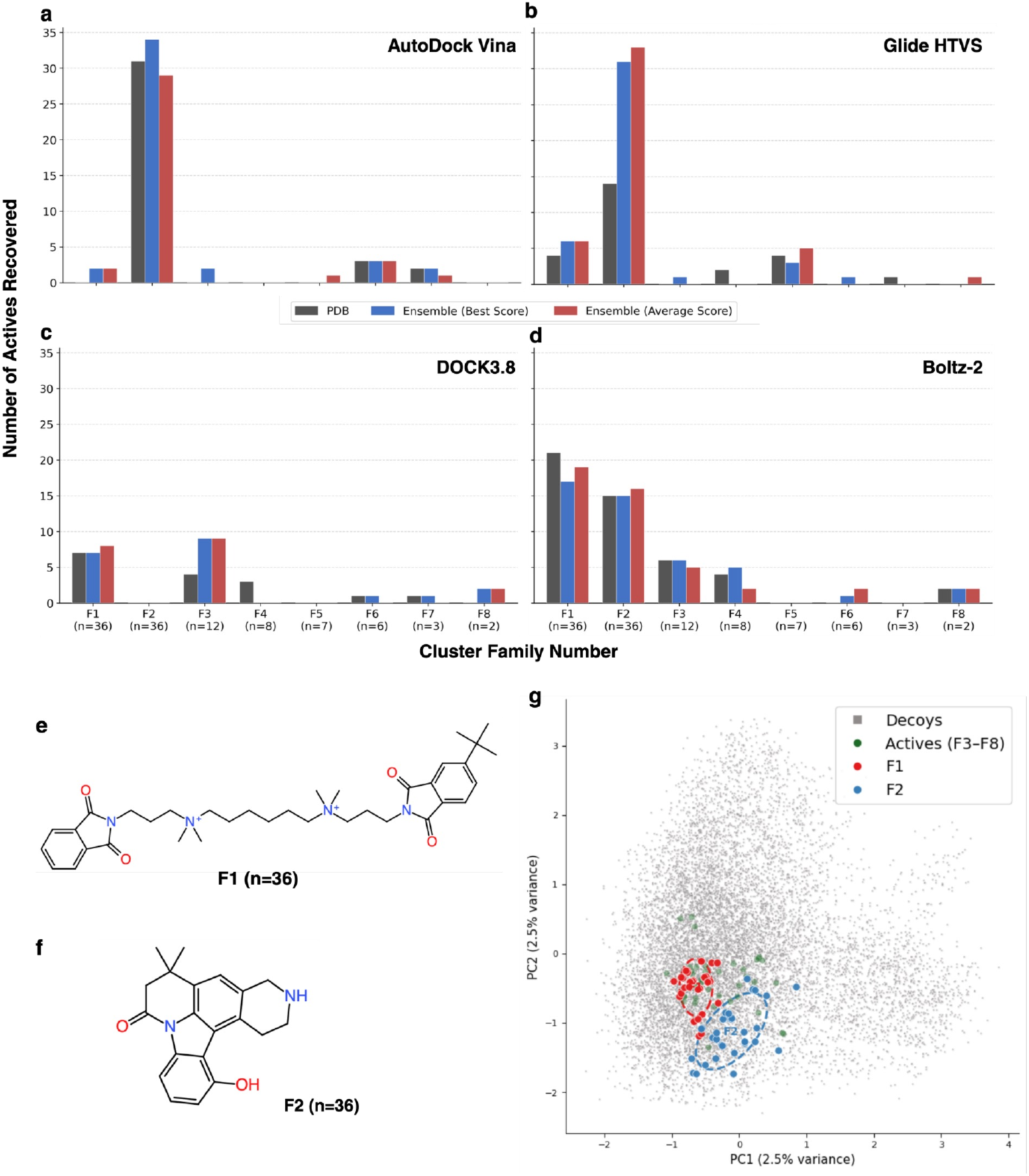
Chemical Similarity and Clustering of M2R Actives. Clustering (RDKit circular fingerprints; 2048 bits) of the top 5% of M2R actives recovered across all docking protocols (n=116). (a-d) The top 8 families, based on cluster size, and the number of actives recovered from each cluster family by (a) AutoDock Vina, (b) Glide HTVS, (c) DOCK3.8, and (d) Boltz-2. (e-f) Representative scaffolds from the top two most populated clusters, F1 and F2, respectively. (g) For chemical space visualization, extended-connectivity fingerprints (ECFP4) (radius = 2, 2048 bits) were computed for all M2R actives (n = 230) and decoys (n = 11,500) and projected onto two principal components fit on the combined active + decoy fingerprint matrix. Confidence ellipses for F1 (blue) and F2 (red) represent 1.5 standard deviations of the PC1/PC2 distribution within each scaffold family. Decoys are given for background (grey), and the non-F1/2 actives are in green.

### Allosteric Pocket Topology Influences Per-Cluster Docking Utility

Per-cluster analysis of M2R ensemble docking revealed that clusters with greater simulation occupancy and lower reweighted free energy generally yielded improved early enrichment across Glide and Vina (**Figs. 2c, 7b-c**). However, pocket topology introduced additional failure modes displayed by both programs that free energy alone could not anticipate. Most clusters maintained structurally similar allosteric pocket conformations relative to cluster 00, with a few exceptions, such as clusters 05 and 07, which each deviated by more than 2.0 Å (**Fig. 7a**). These clusters demonstrated poor enrichment across both Vina and Glide. Cluster 07 presented a substantially shallower binding surface (**Fig. 7h**), poor SiteScore, and unfavorable hydrophilic-to-hydrophobic balance (**Table 2**), yielding a geometrically challenging pocket that neither program’s search algorithm could productively explore. Cluster 03’s failure was for different reasons: with one of the lowest RMSD values in the ensemble, near-identical pocket volume to the PDB (320.70 vs. 322.08 Å^3^, **Table 2**), and a similarly wide, solvent-exposed surface topology (**Fig. 7e**), cluster 03 effectively recapitulates the same geometric limitations that make the PDB a weak docking template for M2R PAMs. This explains its consistently poor enrichment despite its thermodynamic favorability.

**Figure 7.**
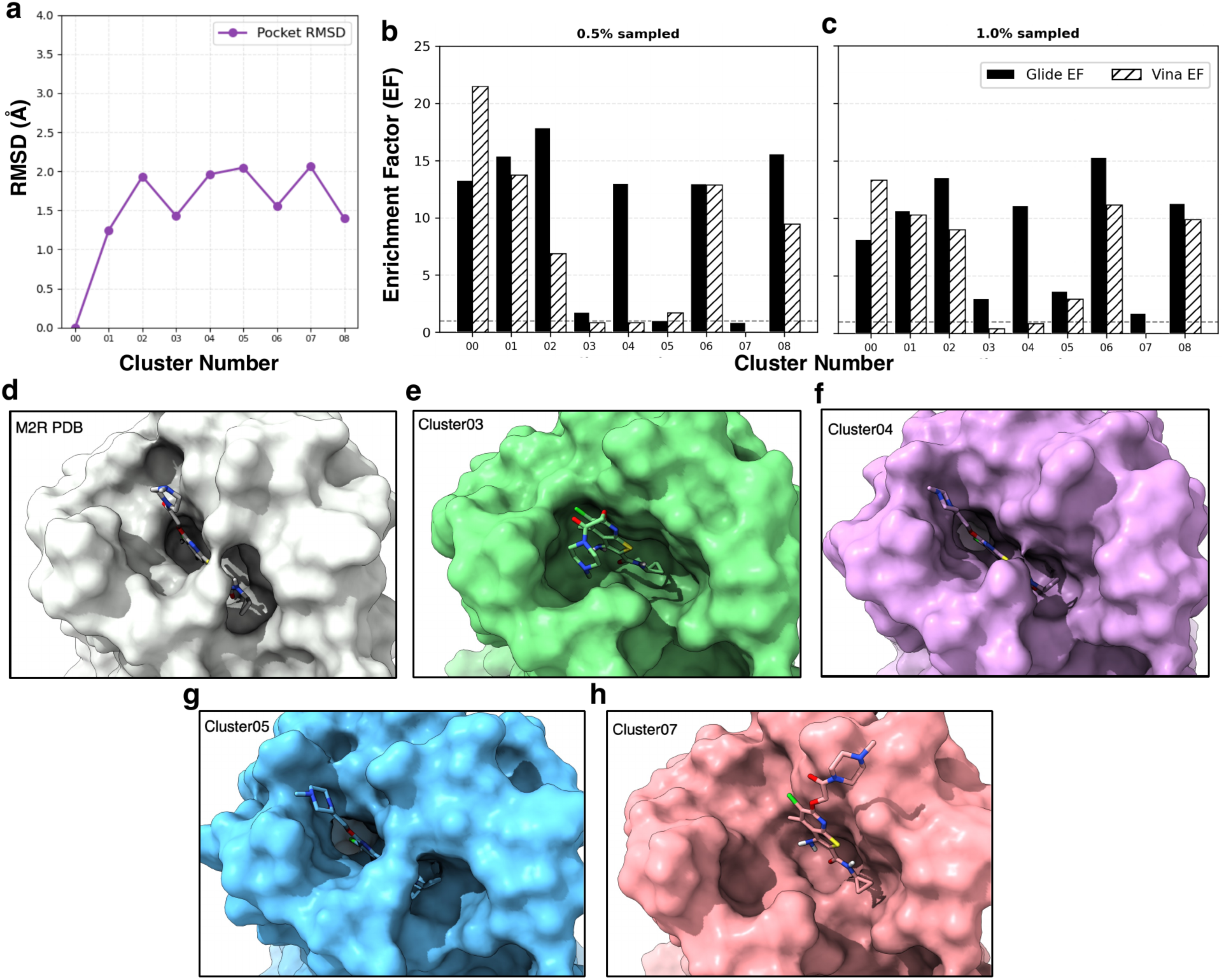
Structural Basis of Per-cluster Performance. (a) RMSD of the allosteric pocket in GaMD clusters of the M2R relative to the cluster 00. Per-cluster EF at (b) 0.5% and (c) 1% sampled for Glide (solid) and Vina (hashed). (d) Surface topology of the M2R PDB compared with select M2R ensemble clusters: (e) cluster03, (f) cluster04, (g) cluster05, and (h) cluster07.

The starkest divergence in docking performance occurred at cluster 04, which is among Glide’s best-performing individual clusters yet simultaneously among Vina’s worst. This compact, highly hydrophobic pocket (volume 93.98 Å^3^, balance 3.663) represents a relevant conformational transition in the allosteric ensemble (**Fig. S1a**), and its inclusion either boosted or diminished ensemble enrichment depending on the program and re-ranking method used (**Fig. 3c–d**). Cluster 04 performance demonstrated that a single structural cluster can simultaneously represent the most productive conformation for one scoring function and the least productive for another, a contingency invisible to any selection criterion based on pocket properties or RMSD alone. Collectively, GaMD-derived ensembles provide M2R allosteric pocket coverage that no single PDB structure can replicate, but productive coverage requires compatibility between individual cluster geometry and the scoring function and search algorithm of the chosen program.

### Diverse Free Energy Landscapes and Pocket Properties Across Structural Ensembles of Different GPCRs

We next evaluated whether our M2R findings generalize across three additional Class A GPCR targets with distinct allosteric mechanisms and pocket geometries: the M4R (extracellular vestibule PAM site, n=2,314 PAMs), CCR2 (intrahelical NAM site, n=16 NAMs), and β2AR (intrahelical NAM site, n=64 NAMs). The same protocol was applied for structure preparation, GaMD structural clustering, compound library curation, and docking, using Glide and Vina, the top two programs identified from M2R benchmarking. GaMD simulations captured meaningful allosteric pocket flexibility across all three targets, with pocket RMSD fluctuating between 1.4 ± 0.5 Å for M4R (**Fig. S7a**), 1.9 ± 0.3 Å for the CCR2 (**Fig. S8a**), and 1.7 ± 0.1 Å for the β2AR (**Fig. S9a**). The free energy landscapes, however, differed substantially across targets: CCR2 clusters 00-03 were frequently revisited throughout the simulations, reflecting a relatively flat multi-basin landscape, whereas M4R and β2AR were dominated by cluster 00 with steep free energy gradients separating the ground state from higher-energy clusters (**Fig. S10**). SiteMap analysis confirmed that ensemble clusters from each target shared druggability properties with their respective PDB structures, displaying good binding site scores (SiteScore >1.0), high hydrophobicity, and hydrogen bond acceptor character across the allosteric pocket region (**Tables S2-S4**), indicating that the GaMD ensembles broadly sampled productive binding-competent conformations.

### M4R Ensemble Docking Favors *BE_min_* Re-ranking Due to Skewed Free Energy Landscape

Among the 12 early enrichment metrics (EF(0.5-1%), EF′(0.5-1%), AUC, and logAUC) computed for M4R, 9 (75%) improved with ensemble docking relative to PDB docking alone (**Table 4**). All 6 Glide metrics improved under *BE_min_*, while only 3 Vina metrics showed improvement with no consistent preference for either ensemble re-ranking method. Glide displayed superior early-ranking precision at M4R relative to Vina (**Fig. 8a,d**), which, despite Vina’s competitive global discrimination (AUC: PDB 72.53%; *BE_min_* 73.38%) (**Fig. 9d**), lacked the early-percentile concentration of actives that would support its prospective screening utility. Notably, the Glide M4R *BE_min_* ensemble model outperformed the *BE_avg_* model at top percentiles, contrasting with Glide M2R docking, where the *BE_avg_* model enrichment improved as additional clusters were incorporated (**Fig. 3c-d**). While the M2R ensemble sampled a broad range of allosteric conformations within a compact free energy range (0-2.83 kcal/mol), the M4R ensemble was dominated by cluster 00 (∼80% of frames) with a steeper free energy gradient across the remaining clusters (0-5.22 kcal/mol, **Fig. S10a**). For M4R, adding cluster 01 alone produced a 9.8-point EF(0.5%) decline under *BE_avg_*, with performance decreasing further with each subsequent cluster (**Fig. S11a**). The large free energy penalties carried by sparsely populated M4R clusters dilute the enrichment signal when averaged across the full ensemble, whereas *BE_min_* naturally recovered the best per-compound score regardless of ensemble composition. This aligns with our findings from M2R benchmarking: *BE_avg_* was most beneficial when the free energy landscape of GaMD receptor structural clusters were flat and well-populated, while *BE_min_* was more robust when sampling was dominated by a single low-energy ground state.

**Figure 8.**
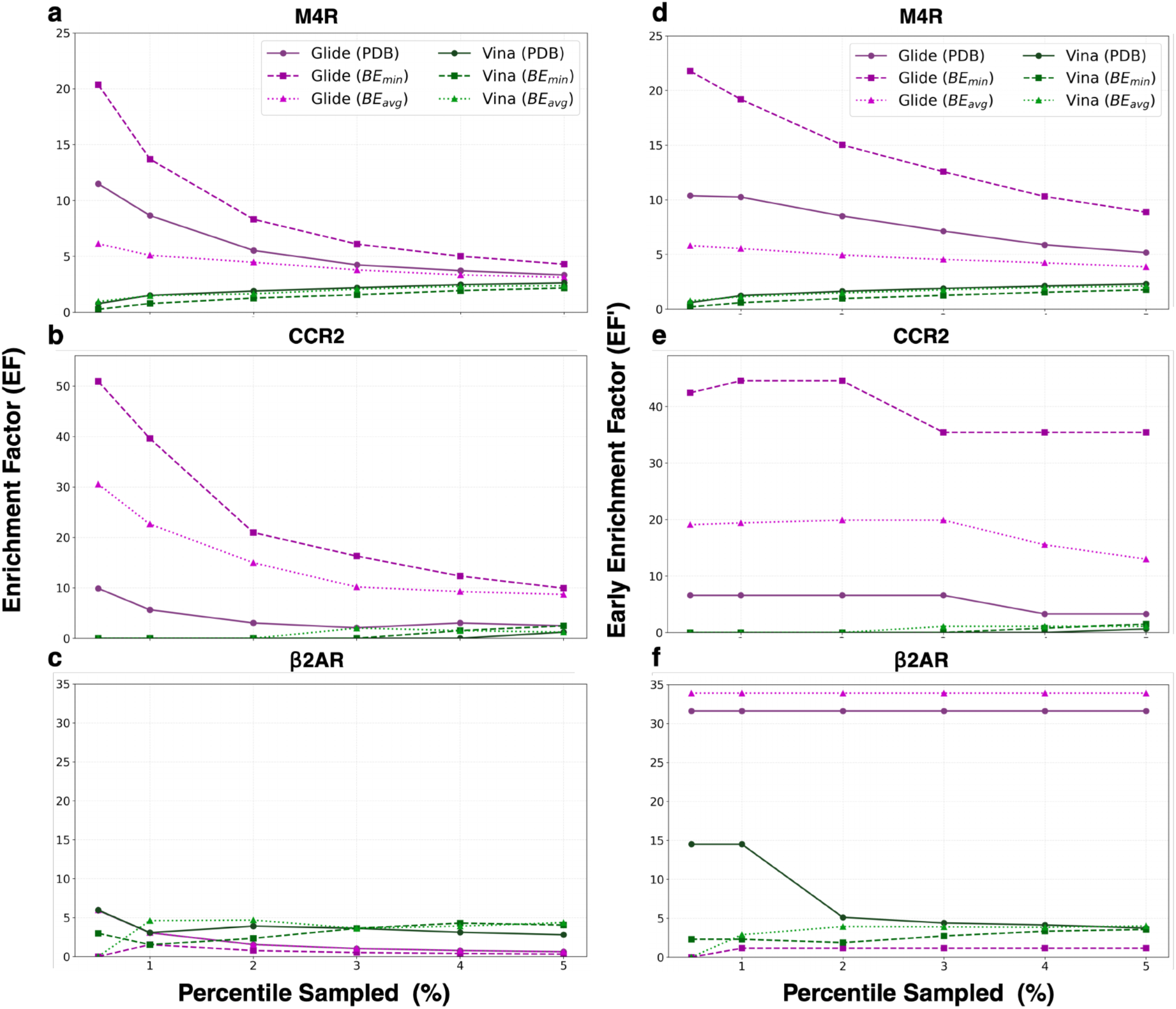
M4R, β2AR, and CCR2 Enrichment Factors. (a-c) Enrichment Factors (EF) and (d-f) Early EF (EF’) for M4R, β2AR, and CCR2, respectively, comparing PDB and ensemble docking performance using Glide (purple) or Vina (green).

**Figure 9.**
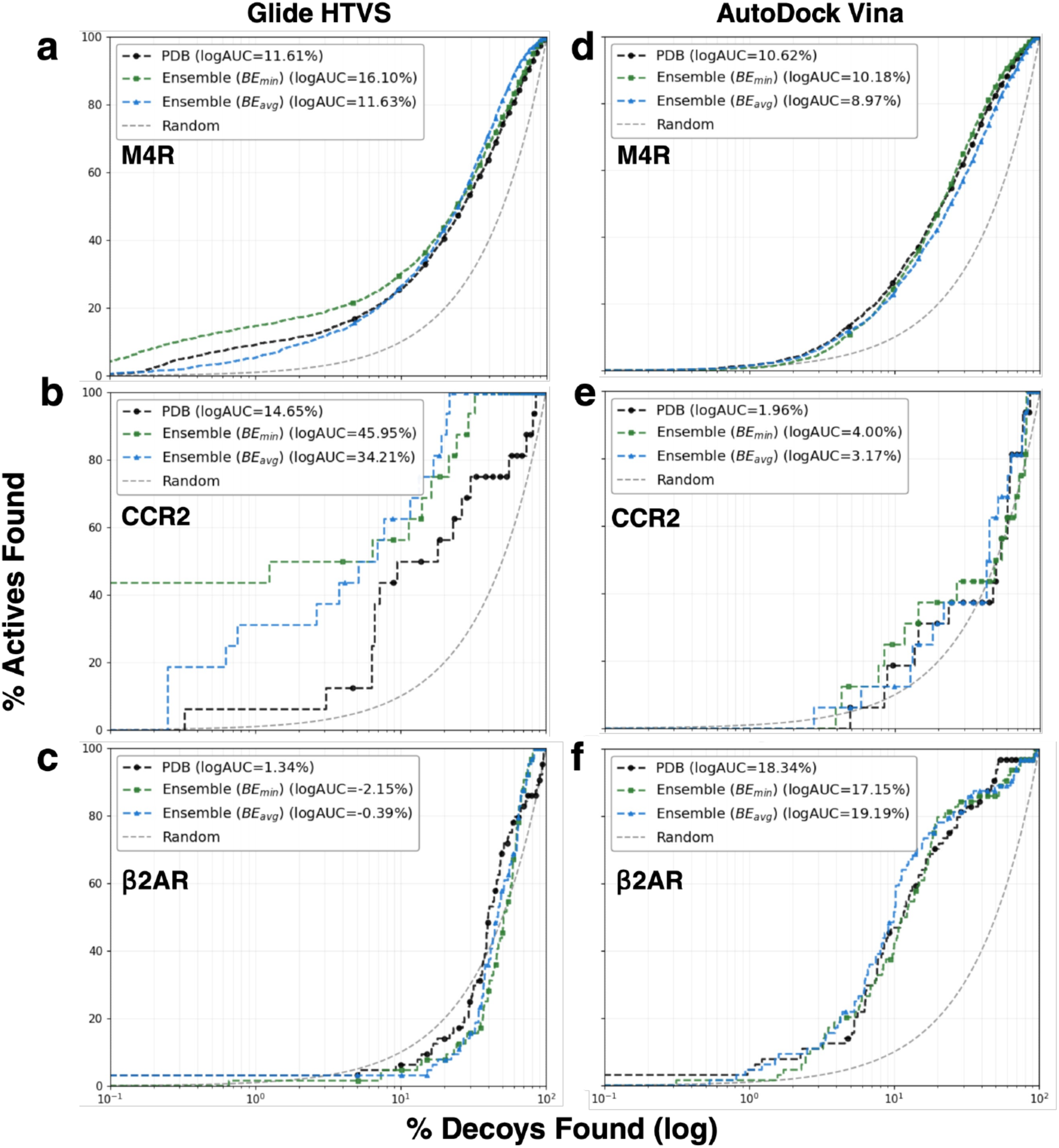
ROC Curves of Allosteric Modulator Docking at M4R, β2AR, and CCR2. Semilogarithmic ROC curves and the (log) Area Under the Curve (AUC) values for PDB (solid black) and ensemble docking (*BE_min_* and *CS_max_* [green, top ensemble docking score]; *BE_avg_* and *CS_avg_* [blue, average ensemble docking score])of allosteric modulators to M4R, CCR2, and β2AR structures, using (a-c) Glide or (d-f) Vina, respectively.

**Table 4.**
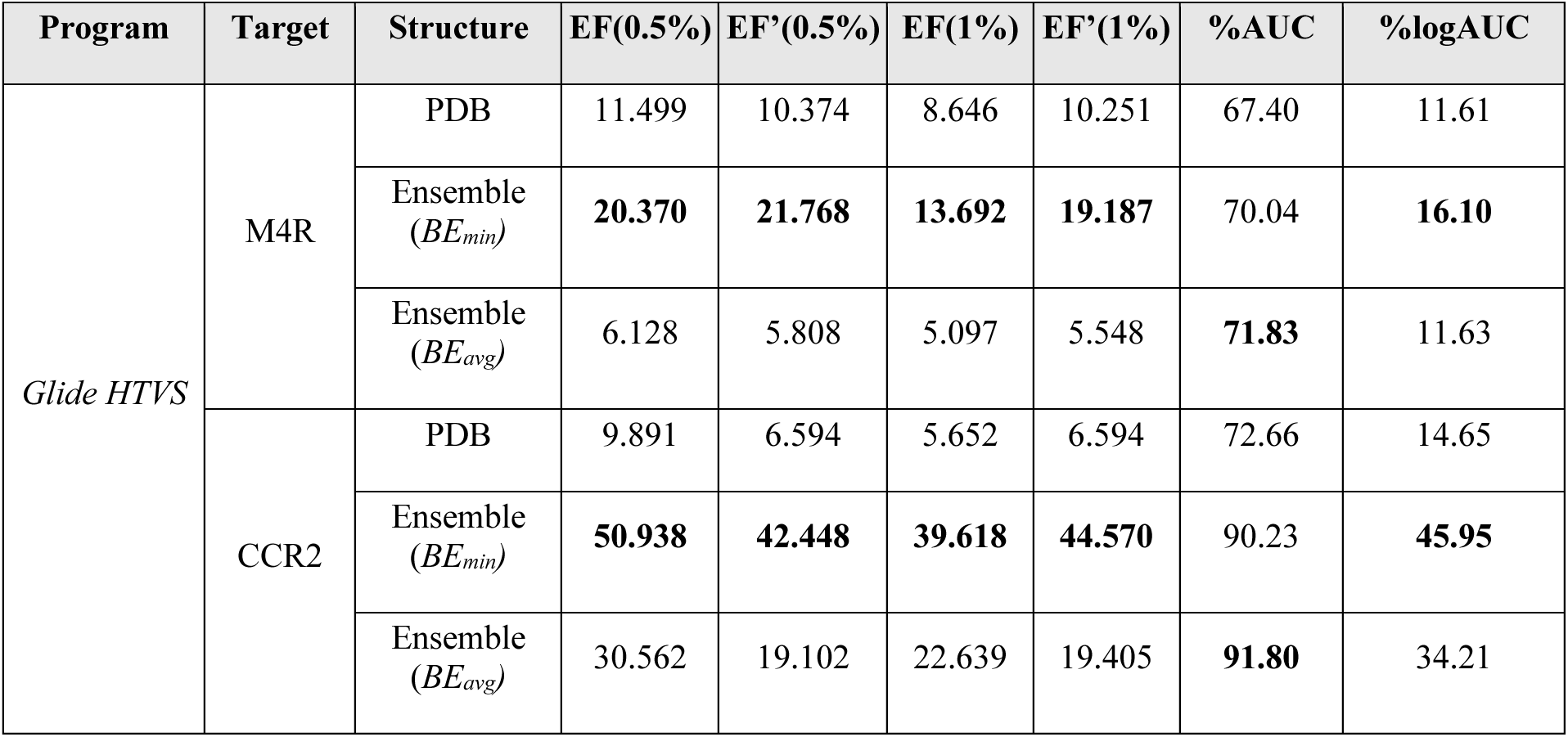
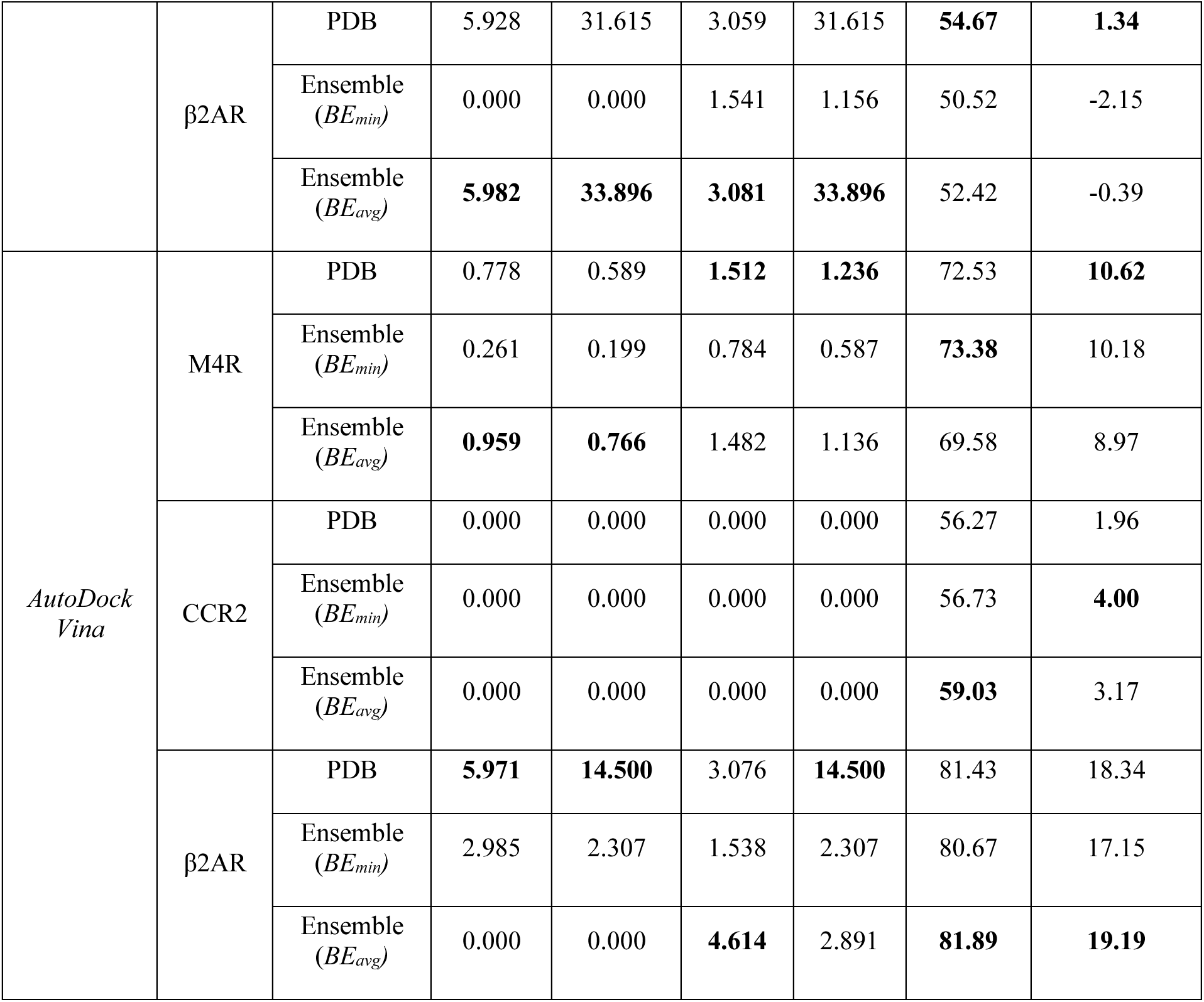
Enrichment Factors and AUC Values from PDB and Ensemble Docking of M4R, CCR2, and β2AR Allosteric Modulators Using Glide and Vina.

| Program | Target | Structure | EF(0.5%) | EF'(0.5%) | EF(1%) | EF'(1%) | %AUC | %logAUC |
| --- | --- | --- | --- | --- | --- | --- | --- | --- |
| Glide HTVS | M4R | PDB | 11.499 | 10.374 | 8.646 | 10.251 | 67.40 | 11.61 |
| | | Ensemble ($BE_{min}$) | **20.370** | **21.768** | **13.692** | **19.187** | 70.04 | **16.10** |
| | | Ensemble ($BE_{avg}$) | 6.128 | 5.808 | 5.097 | 5.548 | **71.83** | 11.63 |
|  | CCR2 | PDB | 9.891 | 6.594 | 5.652 | 6.594 | 72.66 | 14.65 |
| | | Ensemble ($BE_{min}$) | **50.938** | **42.448** | **39.618** | **44.570** | 90.23 | **45.95** |
| | | Ensemble ($BE_{avg}$) | 30.562 | 19.102 | 22.639 | 19.405 | **91.80** | 34.21 |
| | β2AR | PDB | 5.928 | 31.615 | 3.059 | 31.615 | **54.67** | **1.34** |
| | | Ensemble<br>($BE_{min}$) | 0.000 | 0.000 | 1.541 | 1.156 | 50.52 | -2.15 |
| | | Ensemble<br>($BE_{avg}$) | **5.982** | **33.896** | **3.081** | **33.896** | 52.42 | -0.39 |
| *AutoDock*<br>*Vina* | M4R | PDB | 0.778 | 0.589 | **1.512** | **1.236** | 72.53 | **10.62** |
| | | Ensemble<br>($BE_{min}$) | 0.261 | 0.199 | 0.784 | 0.587 | **73.38** | 10.18 |
| | | Ensemble<br>($BE_{avg}$) | **0.959** | **0.766** | 1.482 | 1.136 | 69.58 | 8.97 |
|  | CCR2 | PDB | 0.000 | 0.000 | 0.000 | 0.000 | 56.27 | 1.96 |
| | | Ensemble<br>($BE_{min}$) | 0.000 | 0.000 | 0.000 | 0.000 | 56.73 | **4.00** |
| | | Ensemble<br>($BE_{avg}$) | 0.000 | 0.000 | 0.000 | 0.000 | **59.03** | 3.17 |
| | β2AR | PDB | **5.971** | **14.500** | 3.076 | **14.500** | 81.43 | 18.34 |
| | | Ensemble<br>($BE_{min}$) | 2.985 | 2.307 | 1.538 | 2.307 | 80.67 | 17.15 |
| | | Ensemble<br>($BE_{avg}$) | 0.000 | 0.000 | **4.614** | 2.891 | **81.89** | **19.19** |

### Glide Achieves Near-Perfect Early Enrichment at CCR2

CCR2 ensemble docking with Glide yielded the most substantial enrichment gains relative to the PDB structure (**Fig. 8b, 9b**; **Table 4**). The *BE_min_* model produced over 40-and 36-point improvements in EF and EF’, respectively, identifying almost exclusively actives at the earliest sampled percentiles. This high-level performance was strikingly achieved by screening against cluster 00 alone, with subsequent clusters offering marginal additional benefit (**Fig. S12a-f**). This one GaMD-derived conformation offers a substantially more productive screening starting point than the experimental PDB structure itself. This may offer an efficient and cost-effective alternative to full ensemble screening. Vina’s performance at CCR2 presented a stark contrast. Both the PDB and ensemble models returned the lowest enrichment values of any evaluated target (**Fig. 8e,9e**), with no actives recovered below 3% sampled under any protocol and PDB docking failing to return any actives until 5% (**Table 4**). The severity of this divergence between Glide and Vina at a single target may reflect scoring function incompatibilities with the CCR2 allosteric pocket in tandem with the physicochemical character of the NAM library, rather than a deficiency in the ensemble structures themselves.

### β2AR Ensemble Docking Yields Modest Gains, with Rare High-Energy Clusters Contributing to NAM Enrichment

β2AR ensemble docking, in contrast to the high-performing M2R, M4R, and CCR2, yielded only modest performance gains, with only 7 of the 12 (58.33%) metrics improving relative to the PDB across Glide and Vina (**Table 4**). Glide *BE_avg_* enrichment was almost equivalent to the PDB (**Fig. 8c**), and AUC and logAUC values were marginally worse (**Fig. 9c**). This is the only instance across all four targets where ensemble docking failed to improve upon random-baseline performance at the earliest percentiles. Similarly, Vina showed slight ensemble success, with EF(1%), AUC, and logAUC marginally improved (**Table 4**). However, Vina’s global discrimination at β2AR was much stronger than Glide’s, with AUC values reaching ∼81% and higher logAUC values across both PDB and ensemble models (**Fig. 9f**). In contrast to M4R and CCR2, screening against cluster 00 alone did not yield the best enrichment for either program despite it being the most populated (∼88%) of the ensemble (**Fig. S10c**). Instead, β2AR ensemble enrichment improved incrementally with additional clusters: Glide plateaued after cluster 01, while Vina continued to improve through clusters 02-08 at later percentiles (2-5%), despite these representing increasingly rare, high-energy allosteric states (**Fig. S13**). This decoupling of population dominance from docking utility suggests that the thermodynamically preferred ground-state conformation is not the most geometrically complementary to the β2AR NAM series. β2AR NAMs may preferentially stabilize minor allosteric pocket states that are rarely visited in the GaMD simulation yet critical to NAM recognition.

## DISCUSSION AND CONCLUSIONS

GaMD-derived ensemble docking improved early AM enrichment relative to single PDB structure docking under at least one program and re-ranking strategy across all four GPCR targets (Fig. 10). These results highlight the broader utility and success of receptor structural ensembles for GPCR allosteric VS. Glide HTVS was the only program to improve enrichment at every target evaluated (Fig. 10a-db), with the largest absolute gains at CCR2 (>40-point EF improvement under *BE_min_*, with nearly all actives placed in the top 0.5% of ranked compounds) driven by cluster 00 alone. This trend was continued at M4R where *BE_min_* improved all six early enrichment metrics. Vina, by contrast, showed greater target-dependent diversity: it was competitive with Glide at M2R, offered strong global discrimination at β2AR (AUC ∼81%), but failed to recover any actives at CCR2. These differences emphasize the risk of committing to a single docking program for GPCR AM VS without prior retrospective validation but demonstrate the benefit of using the GaMD-derived structural ensemble, which generally outperformed the PDB structure across all programs. It is important to note, however, that the magnitude and consistency of these improvements varied substantially by target and program. Ensemble docking success is governed by the same factors that limit PDB docking, e.g., scoring function, conformational search algorithm, and compound library properties, in addition to the conformational free energy landscape of the allosteric pocket.

**Figure 10.**
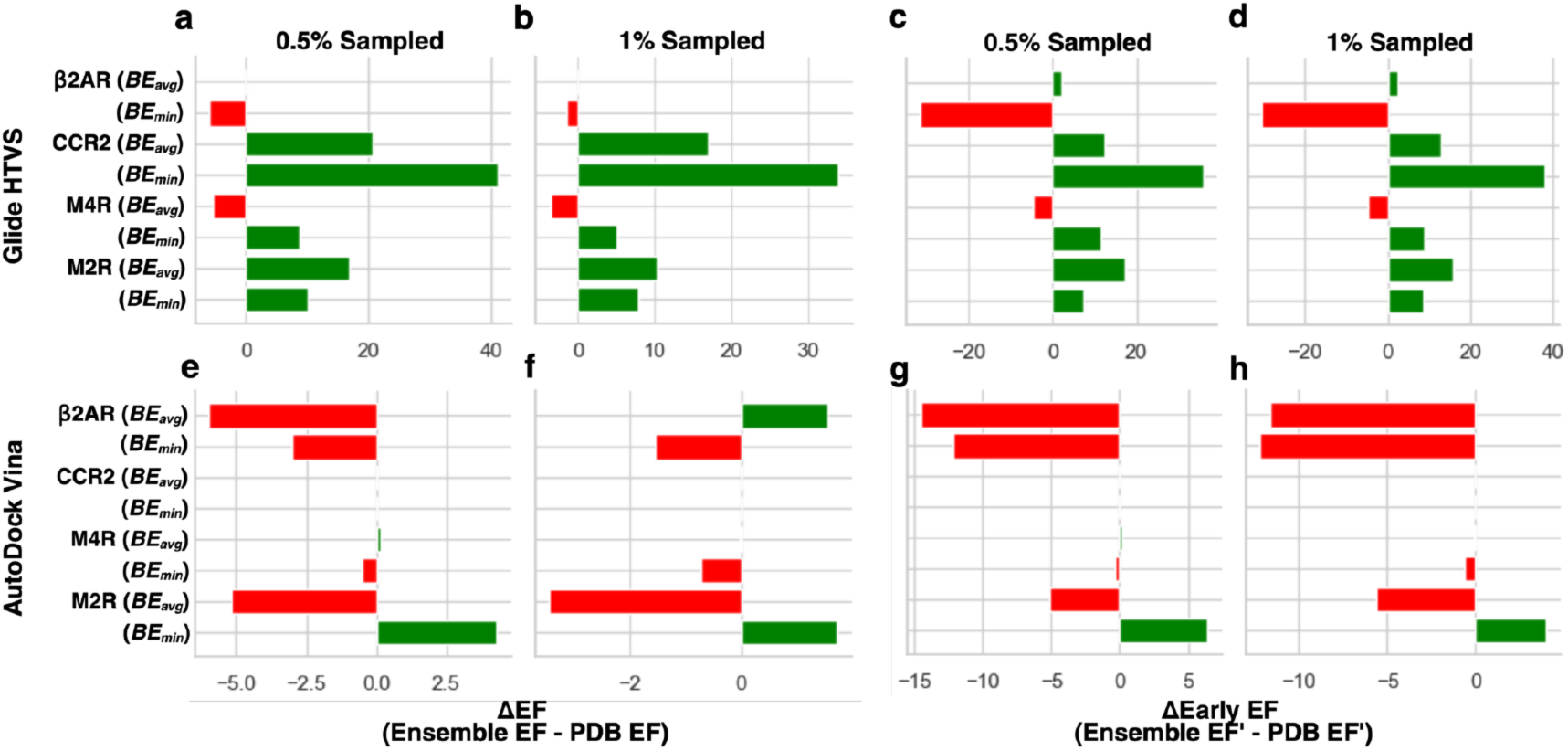
Enrichment Improvement from Using Structural Ensembles for GPCR Allosteric Modulator Docking. The change in magnitude of Glide HTVS (a-b) Enrichment Factors (EF) and (c-d) Early EF (EF’) at 0.5% and 1% sampled from using structural ensembles as opposed to the PDB across all targets (M2R, M4R, CCR2, and B2AR). The change in magnitude of AutoDock Vina (e-f) EF and (g-h) EF’ at 0.5% and 1% sampled from using structural ensembles as opposed to the PDB across all targets (M2R, M4R, CCR2, and B2AR). The more positive the value, the greater the relative improvement ensemble docking provides compared to PDB docking (green, right-side projecting bars). Negative values indicate that ensemble docking yielded worse enrichment than PDB docking (red, left-side projecting bars). Values of zero indicate identical docking performance.

The GaMD reweighted free energy landscape of each target directly influenced which ensemble re-ranking strategy produced the best early enrichment (**Figs. 2c, S10**). At M2R, where clusters 00-03 collectively accounted for ∼87% of sampled frames within a free energy range of 0-2.83 kcal/mol, *BE_avg_* enrichment improved steadily with ensemble size for Glide (**Fig. 3c-d**). At M4R and CCR2 (**Figs. 8a-e**), where a single cluster dominated sampling (∼80% and ∼64% of frames, respectively) and free energy rose steeply thereafter, *BE_min_* consistently outperformed *BE_avg_*. Here, contributions from sparsely populated high-energy clusters diluted the strong enrichment signal concentrated in the ground state. β2AR, despite having the most population-skewed landscape of all targets (∼88% in cluster 00), did not follow this pattern. This target’s enrichment accumulated incrementally across higher-energy clusters rather than being driven by cluster 00 alone (**Fig. S13**), suggesting that β2AR NAMs may preferentially recognize minor allosteric pocket states inaccessible to PDB docking. When the GaMD free energy landscape is flat and multi-populated, *BE_avg_* is the preferred re-ranking strategy; when sampling is dominated by a single low-energy cluster, *BE_min_* is more robust.

Despite differences in early recovery rates between *BE_min_* and *BE_avg_*, almost all M2R actives recovered at the 3% percentile were identified by both re-ranking methods, regardless of program (**Fig. 5**). At these later percentiles, the two ensemble strategies are complementary rather than mutually exclusive, motivating a consensus scoring approach for prospective campaigns. By taking the union of hits ranked highly by both *BE_min_* and *BE_avg_* at the earliest percentiles, you maximize the chemical diversity of the hits. The importance of this recommendation is underscored by the substantial program-specific scaffold recovery biases identified across all protocols (**Fig. 6**). Clustering of M2R actives recovered at 0.5-5% sampled (n = 116) yielded 14 distinct chemical families (F1-F14), with Glide and Vina strongly enriching the high-similarity F2 scaffold family (iSIM=0.853) (**Fig. 6a-b**), while DOCK3.8 and Boltz-2 preferentially recovered the more diverse F1 family (iSIM=0.658) (**Fig. 6c-d**). Boltz-2 uniquely captured the broadest scaffold distribution across all four families with near-equal representation across PDB and ensemble protocols — suggesting its affinity prediction module samples chemically distinct active space from conventional scoring functions, even at lower early enrichment. GaMD ensemble docking recovered actives across scaffold families that PDB-only screening would have missed entirely, demonstrating that conformational diversity in the ensemble translates directly into chemical diversity in recovered hits. This is a key advantage for ensemble-based prospective campaigns where scaffold novelty is often a top priority.

Per-cluster analysis of the M2R ensemble further revealed that not all ensemble structures contribute equally to screening success (**Fig. 7b-c**). Clusters with high pocket RMSD and unfavorable SiteMap properties, such as cluster 07, provided little enrichment benefit regardless of program, while individual cluster success varied substantially by program. M2R cluster 04 exemplifies this most: a compact, highly hydrophobic pocket that ranked among Glide’s best-performing individual frames yet simultaneously among Vina’s worst, with its inclusion either improving or dampening *BE_avg_* ensemble enrichment based entirely on the program used. As a result, per-cluster retrospective benchmarking is an essential step in ensemble protocol optimization, enabling informed decisions about which conformations to include or exclude before committing to a prospective screen. Despite strict template-forcing with a GaMD-derived M2R ensemble, Boltz-2 predictions converged toward the cryo-EM ground state regardless of the template provided, rendering the allosteric pocket conformational diversity largely irrelevant to compound ranking (**Figs. 3** and **S5**). As DL affinity prediction tools continue to mature, incorporating richer allosteric training data and more flexible conformational conditioning will be critical to making these methods competitive with physics-and empirical-based ensemble docking for GPCR AM discovery.

The β2AR represented the most limited ensemble docking improvement across all targets (**Fig. 10**), and the reason for its performance divergence is multifactorial. The weak Glide ensemble performance contrasted with Vina’s competitive early enrichment across both PDB and ensemble models (**Figs. 8f, 9f**), suggesting a program-specific incompatibility with the β2AR NAM chemical series rather than a fundamental failure of the ensemble approach. The small active set size (n=64) and the physicochemical character of the ASD2023 β2AR NAMs likely compounded this difficulty. Expanding the β2AR NAM dataset with experimentally confirmed inactives and evaluating alternative inactive-state structural models would help disentangle scoring function incompatibility from dataset composition effects and represent a natural next step for establishing robust ensemble docking protocols for this target.

This study established GaMD-derived ensemble docking as a broadly beneficial but target-and program-dependent strategy for GPCR AM VS. Here, we provided the first systematic evaluation of GPCR AM enrichment across both conventional docking programs as well as Boltz-2 using both PDB and ensemble structures. Practically, we recommend performing retrospective benchmarking using high-accuracy physics-based or empirical methods tailored to your target, as program-target compatibility varies substantially. In addition, it is imperative to analyze the GaMD free energy landscape of the GPCR allosteric pocket to select between *BE_min_* and *BE_avg_* re-ranking strategies where skewed, population-dominated landscapes favor *BE_min_*, while flat, more evenly distributed landscapes favor *BE_avg_*. We also note that applying a consensus scoring strategy combining both *BE_min_* and *BE_avg_* is ideal to maximize the chemical diversity of recovered hits at the earliest percentiles. Further, testing cluster 00 alone seems to be a worthy effort for a computationally efficient alternative to full ensemble screening. Finally, our work and prior works^71,79^ suggest that, in its current form, Boltz-2 should be used as a complementary rather than primary screening tool for structural prediction and chemical space exploration tasks. Enhanced sampling methods, DL co-folding tools, and allosteric structural databases will continue to develop, and the framework established here provides a rigorous and reproducible foundation for integrating these advances into next-generation GPCR AM discovery pipelines.

## ASSOCIATED CONTENT

### Data Availability Statement

All docking scores, GaMD-derived structural ensembles, compound libraries, and analysis scripts used in this study are openly available at: https://github.com/tylerdt1/gpcr-am-ensemble-docking.git.

### Supporting Information

M2R, M4R, CCR2, and β2AR simulation trajectories and clustering statistics; Glide/Vina/DOCK3.8/Boltz-2 M2R per-cluster enrichment results (0.5-5%); clustering summary of M2R early percentile actives; M4R, CCR2, and β2AR GaMD reweighted free energy values and ensemble cluster populations; Glide/Vina M4R, CCR2, and β2AR per-cluster enrichment results (0.5-5%); Glide/Vina/DOCK3.8/Boltz-2 M2R early enrichment statistics (EF, EF’, logAUC, AUC) from 0.5-5% sampled; SiteMap binding pocket predictions for M2R, M4R, CCR2, and β2AR PDB and ensemble structures (PDF)

## AUTHOR INFORMATION

### Author Contributions

T.D.T. performed data curation, formal analysis, investigation, visualization, writing original draft, review, and editing; Y.M. led project administration, supervision, conceptualization, resources, validation, reviewing, and editing the original draft. All authors have approved the final version of the manuscript.

### Notes

The authors declare no competing financial interest.

## Acknowledgements

We thank past and present Miao Lab members, particularly Jinan Wang, for aiding in the initial data procurement process, Victor Adediwura, and Keya Joshi, for their valuable discussions and input. This work used supercomputing resources with allocation award TG-MCB180049 through the Extreme Science and Engineering Discovery Environment ACCESS, which is supported by National Science Foundation grant number ACI-1548562, project M2874 through the National Energy Research Scientific Computing Center (NERSC), which is a U.S. Department of Energy Office of Science User Facility operated under Contract No. DE-AC02-05CH11231, and Research Computing at the University of North Carolina-Chapel Hill. This work was supported by the National Institutes of Health (R35GM163770) and the startup funding project 27110 at the University of North Carolina-Chapel Hill.

