## Supporting information for "Benchmarking Docking Protocols for GPCR Allosteric Modulators"

Benchmarking Docking Protocols for GPCR Allosteric Modulators – Supporting Information

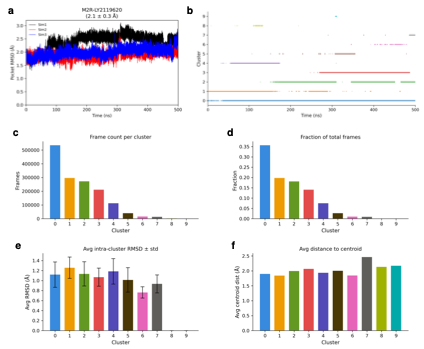

**Figure S1.** M2R Simulation Clustering. (a) Allosteric pocket RMSD fluctuations relative to the PDB over time (3 × 500 ns). (b) CPPTRAJ cluster assignment based on allosteric pocket residues over time. (c) The number of frames allocated to each cluster. (d) The fraction of total frames allocated to each cluster. (e) Intra-cluster RMSD values (average and standard deviation) for each cluster in the ensemble. (f) The average distance from the cluster centroid for each cluster.

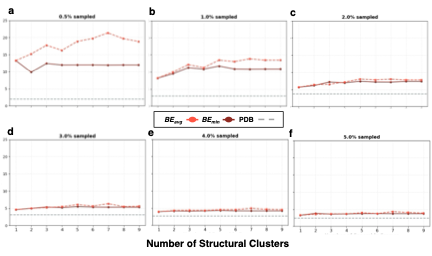

**Figure S2.** Glide Enrichment Results (0.5-5%) from M2R Allosteric Modulator Docking by the Number of Structural Clusters Used for Calculations. (a-f) 0.5-5% sampled EF, respectively, from PDB and ensemble (*BE_min_*, *BE_avg_*) docking.

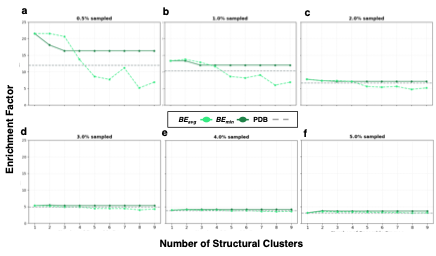

**Figure S3.** Vina Enrichment Results (0.5-5%) from M2R Allosteric Modulator Docking by the Number of Structural Clusters Used for Calculations. (a-f) 0.5-5% sampled EF, respectively, from PDB and ensemble (*BE_min_*, *BE_avg_*) docking.

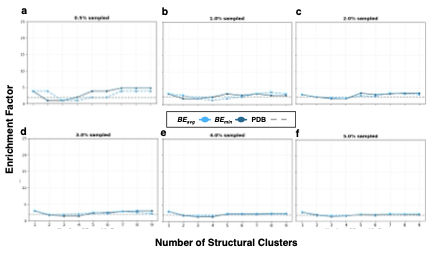

**Figure S4.** DOCK3.8 Enrichment Results (0.5-5%) from M2R Allosteric Modulator Docking by the Number of Structural Clusters Used for Calculations. (a-f) 0.5-5% sampled EF, respectively, from PDB and ensemble (*BE_min_*, *BE_avg_*) docking.

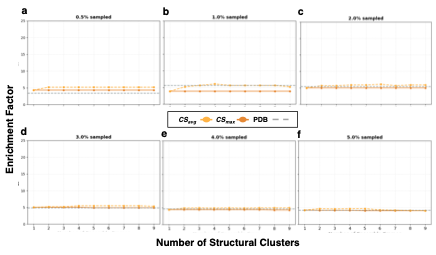

**Figure S5.** Boltz-2 Enrichment Results (0.5-5%) from M2R Allosteric Modulator Docking by the Number of Structural Clusters Used for Calculations. (a-f) 0.5-5% sampled EF, respectively, from PDB and ensemble (*CS_max_*, *CS_avg_*) docking.

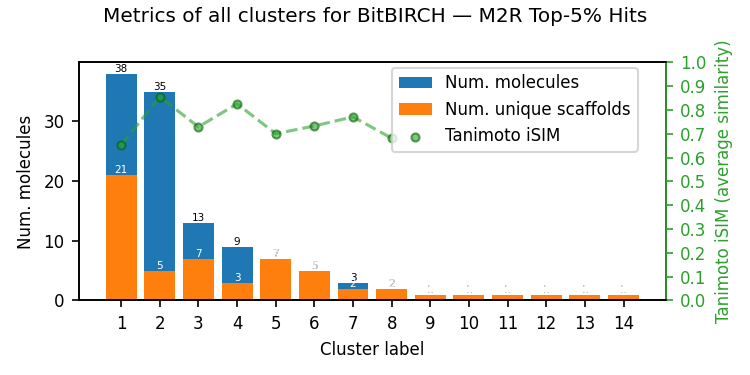

**Figure S6.** BitBirch Clustering Summary of M2R Top Percentile (0.5-5%) Actives. The average Tanimoto similarity (iSIM) is given for each of the eight non-singleton clusters.

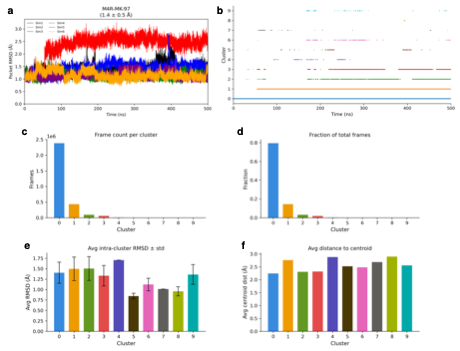

**Figure S7.** M4R Simulation Clustering. (a) Allosteric pocket RMSD fluctuations relative to the PDB over time (6 × 500 ns). (b) CPPTRAJ cluster assignment based on allosteric pocket residues over time. (c) The number of frames allocated to each cluster. (d) The fraction of total frames allocated to each cluster. (e) Intra-cluster RMSD values (average and standard deviation) for each cluster in the ensemble. (f) The average distance from the cluster centroid for each cluster.

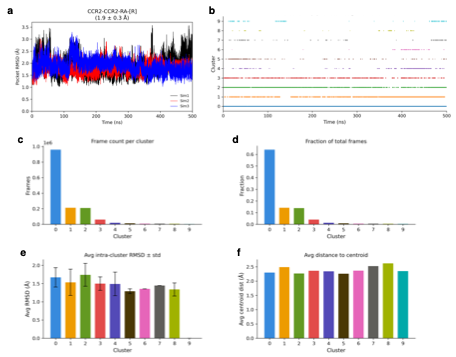

**Figure S8.** CCR2 Simulation Clustering. (a) Allosteric pocket RMSD fluctuations relative to the PDB over time (500ns). (b) CPPTRAJ cluster assignment based on allosteric pocket residues over time. (c) The number of frames allocated to each cluster. (d) The fraction of total frames allocated to each cluster. (e) Intra-cluster RMSD values (average and standard deviation) for each cluster in the ensemble. (f) The average distance from the cluster centroid for each cluster.

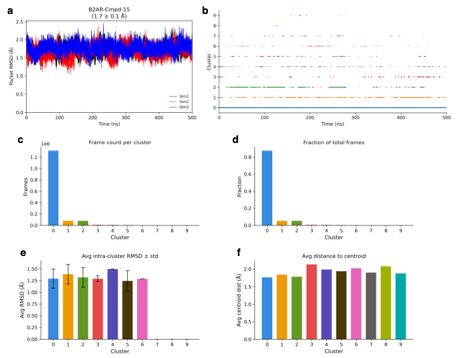

**Figure S9.** β2AR Simulation Clustering. (a) Allosteric pocket RMSD fluctuations relative to the PDB over time (500ns). (b) CPPTRAJ cluster assignment based on allosteric pocket residues over time. (c) The number of frames allocated to each cluster. (d) The fraction of total frames allocated to each cluster. (e) Intra-cluster RMSD values (average and standard deviation) for each cluster in the ensemble. (f) The average distance from the cluster centroid for each cluster.

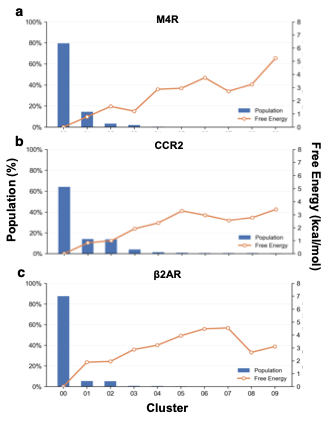

**Figure S10.** GaMD Reweighted Free Energy Distributions and Cluster Population Distributions. Cluster population (%, blue bars; left axis) and the reweighted free energy (kcal/mol, orange line; right axis) from GaMD simulation clustering of (a) M4R, (b) CCR2, and (c) β2AR simulations.

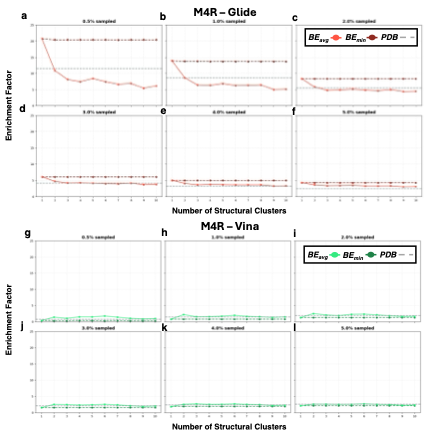

**Figure S11.** Enrichment Results (0.5-5%) from M4R Allosteric Modulator Docking by the Number of Structural Clusters Used for Calculations. 0.5-5% sampled EF, respectively, from PDB and ensemble (*BE_min_*, *BE_avg_*) docking using (a) Glide or (b) Vina.

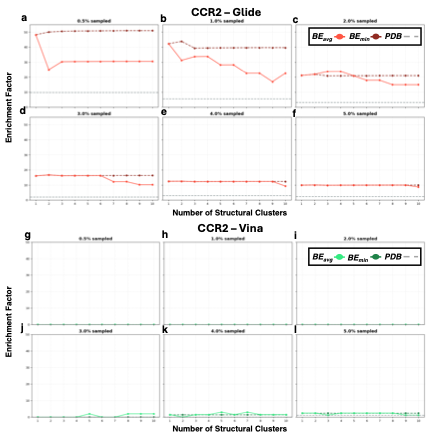

**Figure S12.** Enrichment Results (0.5-5%) from CCR2 Allosteric Modulator Docking by the Number of Structural Clusters Used for Calculations. 0.5-5% sampled EF, respectively, from PDB and ensemble (*BE_min_*, *BE_avg_*) docking using (a) Glide or (b) Vina.

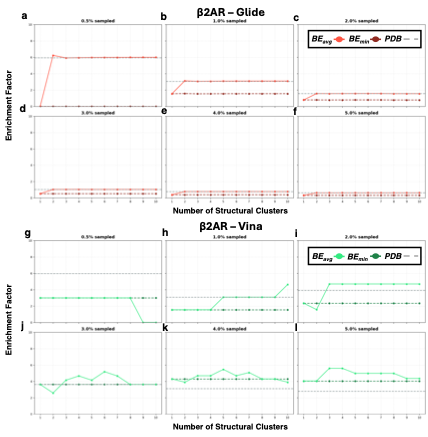

**Figure S13.** Enrichment Results (0.5-5%) from β2AR Allosteric Modulator Docking by the Number of Structural Clusters Used for Calculations. 0.5-5% sampled EF, respectively, from PDB and ensemble (*BE_min_*, *BE_avg_*) docking using (a) Glide or (b) Vina.

**Table S1.** Enrichment Factors from M2R Allosteric Modulator Docking (0.5-5% sampled).

| **Program** | **Target** | **EF(0.5%)** | **EF’(0.5%)** | **EF(1%)** | **EF’(1%)** | **EF(2%)** | **EF’(2%)** | **EF(3%)** | **EF’(3%)** | **EF(4%)** | **EF’(4%)** | **EF(5%)** | **EF’(5%)** |
| --- | --- | --- | --- | --- | --- | --- | --- | --- | --- | --- | --- | --- | --- |
| *Glide HTVS* | PDB | 1.998 | 2.938 | 2.996 | 2.378 | 3.745 | 3.024 | 3.163 | 3.208 | 2.747 | 3.080 | 2.497 | 2.954 |
|  | Ensemble (*BE_min_)* | 11.985 | 10.101 | 10.792 | 10.870 | 7.339 | 10.077 | 5.324 | 9.081 | 4.218 | 8.457 | 3.719 | 7.122 |
|  | Ensemble (*BE_avg_)* | **18.833** | **20.136** | **13.382** | **18.261** | **7.770** | **15.244** | **5.612** | **12.594** | **4.542** | **10.210** | **3.892** | **8.632** |
| *AutoDock Vina* | PDB | 12.030 | 10.218 | 10.312 | 11.548 | 6.688 | 10.300 | 4.898 | 9.257 | 3.892 | 8.322 | 3.115 | **8.322** |
|  | Ensemble (*BE_min_)* | **16.327** | **16.629** | **12.030** | **15.555** | **7.120** | **13.651** | **5.330** | **10.977** | **4.216** | **9.706** | **3.721** | 7.629 |
|  | Ensemble (*BE_avg_)* | 6.874 | 5.128 | 6.874 | 5.901 | 5.178 | 6.187 | 4.321 | 5.506 | 3.676 | 5.102 | 3.202 | 4.742 |
| *DOCK3.8* | PDB | 1.913 | 2.623 | 2.416 | 2.362 | 2.416 | 2.359 | 1.940 | 2.255 | 1.938 | 2.113 | 1.844 | 1.997 |
|  | Ensemble (*BE_min_)* | **4.746** | **4.250** | 2.877 | **3.694** | **3.357** | 3.437 | **3.037** | **3.493** | **2.398** | **3.370** | **2.114** | **3.073** |
|  | Ensemble (*BE_avg_)* | 3.796 | 3.169 | **3.357** | 3.543 | 2.877 | **3.637** | 2.238 | 3.364 | 2.278 | 2.839 | 1.922 | 2.710 |
| *Boltz-2* | PDB | 3.423 | 2.831 | **5.614** | 4.337 | 5.423 | 5.090 | 4.924 | 5.181 | 4.664 | 5.012 | **4.255** | 4.801 |
|  | Ensemble (*CS_max_)* | 4.317 | 4.162 | 3.886 | 3.738 | 4.986 | 4.495 | 4.921 | 4.577 | 4.345 | 4.534 | 4.086 | 4.420 |
|  | Ensemble (*CS_avg_)* | **5.181** | **4.878** | 5.181 | **4.878** | **5.853** | **5.418** | **5.355** | **5.361** | **4.997** | **5.226** | 4.086 | **5.184** |

**Table S2.** SiteMap Binding Pocket Properties for the M4R PDB and Structural Clusters.

| **M4R Structure** | **SiteScore** | **DScore** | **Balance** | **Volume [Å^3^]** | **Hydrophobic** | **Hydrophilic** | **Don/Acc** |
| --- | --- | --- | --- | --- | --- | --- | --- |
| Cluster00 | 1.075 | 1.135 | 1.995 | 342.66 | 1.458 | 0.731 | 0.849 |
| Cluster01 | 1.155 | 1.256 | 6.232 | 359.46 | 2.507 | 0.402 | 0.479 |
| Cluster02 | 1.052 | 1.089 | 0.959 | 377.99 | 0.855 | 0.893 | 0.583 |
| Cluster03 | 1.058 | 1.117 | 1.777 | 418.80 | 1.327 | 0.747 | 0.751 |
| Cluster04 | 1.137 | 1.225 | 3.937 | 349.86 | 1.967 | 0.500 | 0.737 |
| Cluster05 | 1.081 | 1.135 | 1.390 | 396.51 | 1.053 | 0.758 | 1.344 |
| Cluster06 | 1.080 | 1.123 | 1.510 | 423.60 | 1.265 | 0.838 | 0.863 |
| Cluster07 | 1.079 | 1.139 | 1.637 | 325.85 | 1.183 | 0.723 | 0.715 |
| Cluster08 | 1.096 | 1.153 | 1.674 | 417.77 | 1.225 | 0.732 | 1.972 |
| Cluster09 | 1.073 | 1.125 | 1.446 | 427.38 | 1.126 | 0.779 | 0.904 |
| PDB | 1.077 | 1.127 | 1.464 | 349.52 | 1.158 | 0.791 | 0.610 |

**Table S3.** SiteMap Binding Pocket Properties for the CCR2 PDB and Structural Clusters.

| **CCR2 Structure** | **SiteScore** | **DScore** | **Balance** | **Volume [Å^3^]** | **Hydrophobic** | **Hydrophilic** | **Don/Acc** |
| --- | --- | --- | --- | --- | --- | --- | --- |
| Cluster00 | 1.125 | 1.174 | 2.559 | 412.63 | 1.945 | 0.760 | 0.501 |
| Cluster01 | 1.231 | 1.314 | 6.016 | 358.78 | 2.841 | 0.472 | 0.527 |
| Cluster02 | 1.175 | 1.232 | 4.858 | 313.16 | 3.315 | 0.682 | 0.279 |
| Cluster03 | 1.127 | 1.169 | 2.334 | 378.67 | 1.888 | 0.809 | 0.447 |
| Cluster04 | 1.154 | 1.222 | 4.021 | 356.72 | 2.472 | 0.615 | 0.404 |
| Cluster05 | 1.126 | 1.190 | 3.219 | 332.71 | 2.133 | 0.663 | 0.503 |
| Cluster06 | 1.175 | 1.261 | 5.168 | 306.98 | 2.530 | 0.490 | 0.273 |
| Cluster07 | 1.117 | 1.159 | 2.576 | 341.28 | 2.096 | 0.814 | 0.483 |
| Cluster08 | 1.158 | 1.242 | 4.792 | 365.98 | 2.200 | 0.459 | 0.254 |
| Cluster09 | 1.149 | 1.193 | 3.477 | 332.71 | 2.709 | 0.779 | 0.448 |
| PDB | 1.124 | 1.162 | 2.140 | 358.78 | 1.792 | 0.837 | 0.341 |

**Table S4.** SiteMap Binding Pocket Properties for the β2AR PDB and Structural Clusters.

| **β2AR Structure** | **SiteScore** | **DScore** | **Balance** | **Volume [Å^3^]** | **Hydrophobic** | **Hydrophilic** | **Don/Acc** |
| --- | --- | --- | --- | --- | --- | --- | --- |
| Cluster00 | 1.100 | 1.154 | 2.737 | 530.62 | 2.038 | 0.745 | 0.411 |
| Cluster01 | 1.097 | 1.161 | 3.110 | 397.19 | 2.137 | 0.687 | 0.522 |
| Cluster02 | 1.098 | 1.154 | 2.362 | 524.45 | 1.740 | 0.737 | 1.353 |
| Cluster03 | 1.075 | 1.132 | 2.252 | 493.92 | 1.680 | 0.746 | 0.700 |
| Cluster04 | 1.143 | 1.237 | 5.344 | 457.56 | 2.451 | 0.459 | 0.519 |
| Cluster05 | 1.153 | 1.217 | 3.642 | 497.69 | 2.363 | 0.649 | 1.306 |
| Cluster06 | 1.126 | 1.204 | 3.805 | 458.59 | 2.186 | 0.574 | 1.118 |
| Cluster07 | 1.142 | 1.218 | 3.814 | 525.13 | 2.199 | 0.577 | 0.957 |
| Cluster08 | 1.107 | 1.177 | 2.599 | 560.80 | 1.670 | 0.643 | 0.927 |
| Cluster09 | 1.101 | 1.160 | 2.875 | 471.28 | 2.063 | 0.717 | 0.721 |
| PDB | 1.110 | 1.178 | 2.801 | 489.80 | 1.826 | 0.652 | 0.460 |
